# Statistical batch-aware embedded integration, dimension reduction and alignment for spatial transcriptomics

**DOI:** 10.1101/2024.06.10.598190

**Authors:** Yanfang Li, Shihua Zhang

## Abstract

Spatial transcriptomics (ST) technologies provide richer insights into the molecular characteristics of cells by simultaneously measuring gene expression profiles and their relative locations. However, each slice can only contain limited biological variation, and since there are almost always non-negligible batch effects across different slices, integrating numerous slices to account for batch effects and locations is not straightforward. Here, we propose a hierarchical hidden Markov random field model STADIA to reduce batch effects, extract common biological patterns across multiple ST slices, and simultaneously identify spatial domains. We demonstrate the effectiveness of STADIA using five datasets from different species (human and mouse), various organs (brain, skin, and liver), and diverse platforms (10x Visium, ST, and Slice-seqV2). STADIA can capture common tissue structures across multiple slices and preserve slice-specific biological signals. In addition, STADIA outperforms the other three competing methods (PRECAST, fastMNN and Harmony) in terms of the balance between batch mixing and spatial domain identification.

## Introduction

Spatial transcriptomics (ST) technologies can measure gene expression profiles and their relative spatial locations, providing new opportunities and challenges for computational biologists. Various methods have been developed to analyze ST data, including spatial domain identification^1–5^, spatially variable gene detection^6–11^, and spatially aware cell type deconvolution^12–15^. However, most of these approaches only focus on individual ST slices, which may limit their utility for multi-slice analysis. Nevertheless, multi-slice integrative analysis is fundamental for the comprehensive exploration of ST data. It reveals hidden patterns and relationships that may remain concealed when focusing solely on individual slices and enables researchers to capture the intricate spatiotemporal dynamics of gene expression across different slices, providing a more holistic perspective of the underlying biological process. Therefore, it is crucial to consider the integration of multiple ST slices by modeling the gene expression and spatial locations carefully.

When merging multiple slices collected under different conditions, laboratories, or experimenters, there are more or less unwanted factors, often referred to as batch effects. If left unadjusted, these batch effects can mask the biological variation of interest, potentially leading to inaccurate results in downstream analysis. To address this issue, researchers have developed several batch effect correction strategies for single-cell RNA sequencing (scRNA-seq), including ComBat^16^, Harmony^17^, fastMNN^18,19^, Scanorama^20^, and Seurat-CCA^21,22^. However, these approaches are primarily tailored for the removal of batch effects in scRNA-seq without considering the relative locations of different cells. Therefore, applying these methods to multiple ST datasets may not yield the desired results.

In addition, batch effect correction and downstream analysis are usually performed separately both for scRNA-seq^16–22^ and ST^23–25^, which may lead to suboptimal results, as it may not effectively account for the interplay between technical variability and biological variations. This limitation is evident, except for several recent approaches such as BUS^26^, BFR.BE^27^, and PRECAST^28^. BUS corrects batch effects and discovers subtypes by integrating the location-and-scale (L/S) adjustment model^16^ with the Gaussian Mixture Model (GMM)^29,30^ for the analysis of microarray data. BFR.BE is a general sparse factor regression model designed for dimension reduction and batch effect correction. Note that neither BUS nor BFR.BE takes into account the relative positions of different cells. PRECAST jointly estimates low-dimensional representations of biological and systematic variation by factor analysis, while simultaneously conducting spatial clustering using GMM with a latent Markov random field Potts^31^ model. However, PRECAST makes certain assumptions, namely (1) that non-cellular biological variation or batch effects are not orthogonal to the biological space, and the projection of batch effects onto the orthogonal complement of the biological space is discarded, and (2) that local neighboring microenvironments are spatially correlated, captured by an intrinsic conditional autoregressive (CAR) model.

To this end, we develop a new ST Analysis tool for multi-slice integration, DImension reduction and Alignment (STADIA). STADIA is a hierarchical hidden Markov random field model, adapting the BUS and BFR.BE algorithms by further accounting for the relative physical positions of different spots/beads, and relaxing the above two assumptions of PRECAST. Extensive experiments on five ST datasets from different species, different organs, and different platforms, and the comparison with three competing methods including PRECAST, fastMNN and Harmony demonstrated the superior performance of STADIA. Notably, STADIA simultaneously corrects batch effects, identifies shared and slice-specific spatial domains across multiple ST slices, and provide insights into the interpretable different variations among distinct slices,including both additive and multiplicative differences, all within a unified framework.

## Materials and Methods

### Overview of STADIA

STADIA takes the pre-processed gene expression profiles and their corresponding spatial coordinates of multiple ST slices as input. STADIA is a hierarchical hidden Markov random field model (HHMRF) consisting of two hidden states: low-dimensional batch-corrected embeddings and spatially-aware cluster assignments (**Fig. 1a**). Specifically, STADIA first performs both linear dimension reduction and batch effect correction using a Bayesian factor regression model with L/S adjustment. Then, STADIA uses the GMM for embedded clustering. Finally, to ensure local consistency of label assignments, STADIA applies the Potts model on an undirected graph, where nodes are spots from all slices and edges are intra-batch KNN pairs using coordinates and inter-batch MNN pairs using gene expression profiles (**Fig. 1b**). STADIA utilizes the expectation-maximization (EM) algorithm for parameter estimation, which iteratively estimates the missing or latent variables from the observed data in the E-step and then updates the parameters to maximize the posterior distribution.

**Fig. 1.**
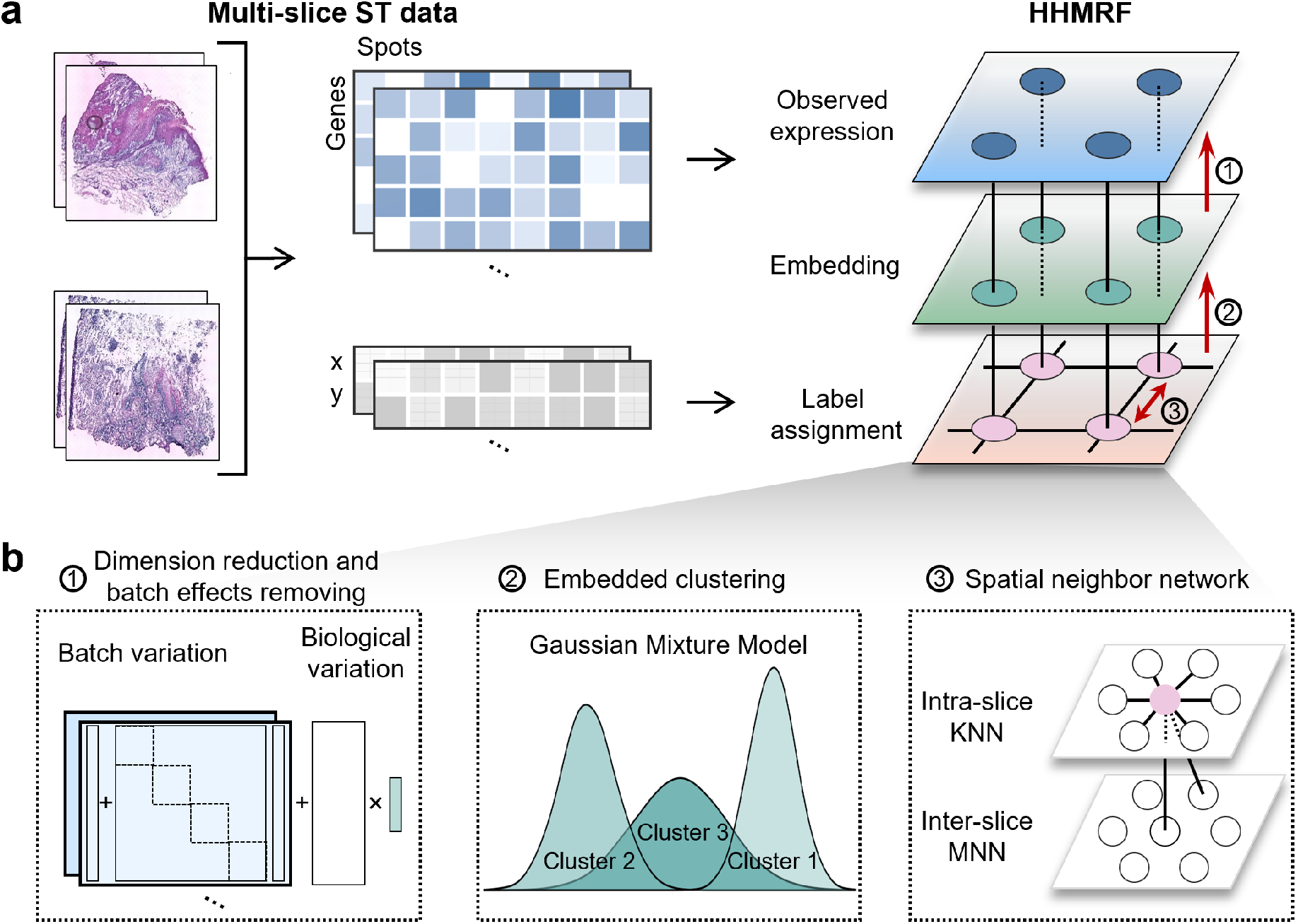
Overview of STADIA. **a**. STADIA is a hierarchical hidden Markov random field model (HHMRF) with multi-slice data as input. **b**. After normal preprocessing on gene expression profiles, STADIA first uses factor analysis and location-and-scale (L/S) adjustment to perform linear dimension reduction and batch effect correction. Then, STADIA uses the Gaussian mixture model to do embedded clustering. Finally, to ensure that the label assignments are locally consistent, STADIA adopts the Markov random field Potts model on the graph, with nodes being spots of all samples and edges being K-Nearest Neighbors (KNN) pairs using coordinates within the batch and mutual nearest neighbors(MNN) pairs using gene expression profiles across batches.

### Data preprocessing

We first performed quality control on all datasets (see **Supplementary Table S1** for details) by filtering out spots expressed in less than 200 genes and genes expressed in less than 20 spots. Then, in each experiment, we selected the top 2000 highly variable genes (HVGs) per slice, ranked them by the number of slices they appear in, and took the top 2000 features as input. Finally, we normalized the raw count data according to the library size and performed a log transformation. We implemented these steps by the Seurat package (v4.3.0)^32^, including the functions `FindVariableFeatures`, `SelectIntegrationFeatures`, `NormalizeData`, and `ScaleData`.

### Construction of the spatial neighborhood graph

We constructed a combined spatial neighborhood graph consisting of all spots from all slices. Two spots within a slice were connected by an edge if the distance between them was among the K-th smallest Euclidean distances. Two spots between any two ST slices were connected if they are mutual nearest neighbors (MNN) based on their gene expression profiles. The undirected graph was represented by an adjacency matrix *A*, where *A*_*ij*_ = 1 if and only if there is an edge between spots *i* and *j*.

### The overall architecture of STADIA

Suppose there are *B* batches. Let ***y***_*bi*_ ∈ ℝ^*p*^ be the observed preprocessed gene expression profile for sample *i* in batch *b* (*i* = 1, *· · ·, n*_*b*_), where *p* is the number of genes measured. Further, denote the true expression levels for sample *i* in batch *b* by ***x***_*bi*_ ∈ ℝ^*p*^ and the vector of batch effects by ***γ***_*b*_ ∈ ℝ^*p*^. Then inspired by the L/S adjustment modeling to remove batch effects for multi-source scRNA-seq^16^, gene expression profiles ***y***_*bi*_ from different batches or slices could be formulated as

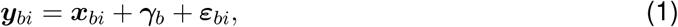

where ***ε***_*bi*_ ∈ ℝ^*p*^ is Gaussian noise with batch-specific diagonal precision matrix **T**_*b*_ that has a distribution equal to 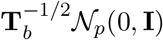, where 𝒩_*p*_(*·*) denotes *p*-dimensional Gaussian distribution. The vectors ***γ***_*b*_ and ***ε***_*bi*_ correct for shifts and proportional changes induced by factors such as variations in instrument calibration and instrument sensitivity, which are referred to as additive and multiplicative batch effects, respectively.

Moreover, the number of genes, *p*, is always ultra-high nowadays due to high-throughput sequencing technology, which is often assumed to lie on a smooth low-dimensional manifold 𝕊^*d*^ (*d* ≪ *p*). Use the linear dimension reduction technique, factor analysis, on ***x***_*bi*_

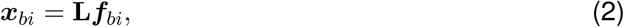

where ***f***_*bi*_ ∈ ℝ^*d*^ is the common factor score for sample *i* in batch *b* and **L** ∈ ℝ^*p×d*^ is the loading matrix shared by all batches. Putting (1) and (2) together, we get the first layer of STADIA

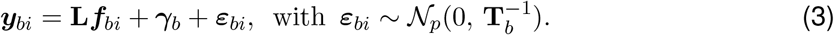

Furthermore, assuming that all samples come from *q* latent clusters, with cluster indicator *c*_*bi*_ for sample *i* in batch *b* and given *c*_*bi*_ = *k* (*k* ∈ [1, …, *q*]), the latent low-dimensional representation ***f***_*bi*_ is normally distributed. More specifically, the second layer of STADIA, which connects the hidden batch-corrected low-dimensional representation and the latent cluster assignments, is a GMM

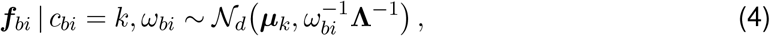

where ***µ***_*k*_ ∈ ℝ^*d*^ is the mean gene expression for the *k*th cluster, *ω*_*bi*_ ∈ ℝ and **Λ** ∈ ℝ^*d×d*^ are sample-specific and common precision, respectively. Note that in such a setting, if the prior distribution of *ω*_*bi*_ is given by the gamma distribution 𝒢 (*ν*_*ω*_*/*2, *ν*_*ω*_*/*2), the marginal distribution of ***f***_*bi*_ by integrating out *ω*_*bi*_ is the multivariate Student-t distribution *t*_*d*_(*ν*_*ω*_, ***µ***_*k*_, **Λ**) with *ν*_*ω*_ degrees of freedom^33^, which is more robust to noise and outliers as specified in BayesSpace^1^.

Finally, since samples close in a physical location within a batch and samples with similar expression across batches tend to have the same biological variations, we use KNN using spatial locations within the slice and MNN using gene expression profiles across slices to construct an undirected graph 𝒢 = (𝒱, ℰ) with nodes 𝒱 representing all spots from all batches. Based on the graph 𝒢, the cluster indicator ***c*** is modeled by the Potts model^31^,

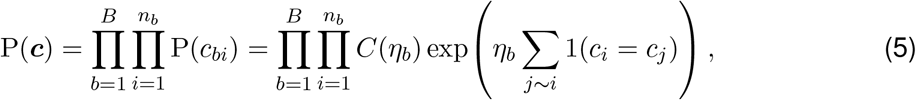

where *C*(*η*_*b*_) is a normalization constant as long as the batch-specific smoothness parameter *η*_*b*_ is fixed beforehand, the notation *j* ∼ *i* denotes all spots connected to spot *i* in the graph 𝒢, representing the neighbors of spot *i*, and 1(*c*_*i*_ = *c*_*j*_) is the indicator function that equals one whenever *c*_*i*_ = *c*_*j*_ and zero otherwise. The Hamiltonian or energy function of the Potts prior (5),

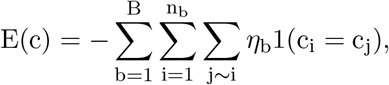

seeks to minimize the discontinuities among neighboring locations by penalizing differences in label assignments ***c*** for adjacent spots in the graph 𝒢. This encourages adjacent spots to have the same cluster or spatial domain.

### Prior formulation

To account for uncertainty in parameter estimates, priors are set for all parameters in this subsection before Bayesian inference (see **Supplementary Note** for details). For the parameters in Eq. (4), weak priors are used to allow for the fact that the data are pretty much nailed down to posterior distributions,

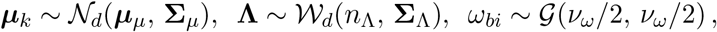

where 𝒲_*d*_(*·*) is the *d*-dimensional Wishart distribution. By default, we set ***µ***_*µ*_ = 0 as a consequence of data centering, and the covariance matrix **Σ**_*µ*_ = 100 × I_*d*_ to down-weight prior to the posterior mean of ***µ***_*k*_. The degree of freedom *n*_Λ_ of the Wishart distribution that satisfies *n*_Λ_ > *d* − 1 determines the certainty of the prior information in the scale matrix, and we set *n*_Λ_ = *d* to provide the least informative specification^34^. The scale matrix is set to **Σ**_Λ_ = 100 *×* I_*d*_. And *ν*_*ω*_ = 2 seems to be successful in avoiding the influence of noise and outliers during segmentation, as set in [35].

Focusing now on Eq. (3), the priors for ***γ***_*b*_ and *t*_*bj*_, the *j*th diagonal of the precision matrix **T**_*b*_, are given respectively by

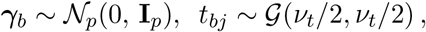

where *ν*_*t*_ = 1. In other words, the additive batch effects ***γ***_*b*_ are considered white noise in our work. For factor-specific gene selection, a traditional Bayesian approach, called spike-and-slab prior, together with the nonlocal product moment (pMOM) prior^36,37^, which satisfies the additional constraint of vanishing probability at point 0, is adopted for **L**,

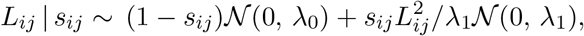

where *s*_*ij*_ ∈ {0, 1}, the variances λ_0_ = 0.015 and λ_1_ = 0.871 are fixed beforehand in all our experiments because 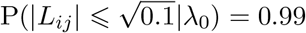 and 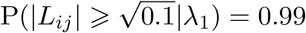. Furthermore, a hierarchical prior is set over the indicator *s*_*ij*_,

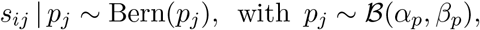

where Bern(*·*) and ℬ (*·*) denote the Bernoulli and Beta distributions, respectively, and with default hyperparameters *α*_*p*_ = *β*_*p*_ = 1. All parameters and their hyperparameters are listed in Table 1, and other hyperparameters can be found in Table 2.

**Table 1.** Parameters with their descriptions and hyperparameters

| Parameter | Description | Hyperparameter |
| --- | --- | --- |
| $\mathbf{L} \in \mathbb{R}^{p \times d}$ | loading matrix | $(\lambda_0, \lambda_1)$ |
| $\boldsymbol{\gamma} \in \mathbb{R}^{p \times B}$ | additive batch-effect vector | |
| $\mathbf{T} \in \mathbb{R}^{p \times B}$ | multiplicative batch effect, precision matrix of error term | $\nu_\tau$ |
| $\boldsymbol{\mu} \in \mathbb{R}^{d \times q}$ | mean matrix of latent representations | $(\boldsymbol{\mu}_\mu, \boldsymbol{\Sigma}_\mu)$ |
| $\boldsymbol{\omega} \in \mathbb{R}^n$ | sample-specific precision of latent representations | $\nu_\omega$ |
| $\boldsymbol{\Lambda} \in \mathbb{R}^{d \times d}$ | shared precision matrix of latent representations | $(n_\Lambda, \boldsymbol{\Sigma}_\Lambda)$ |
| $\mathbf{p} \in \mathbb{R}^p$ | probability in the pMOM prior | $(\alpha_p, \beta_p)$ |

**Table 2.** Other hyperparameters in the model

| Description | Hyperparameter |
| --- | --- |
| smoothing parameter of the Potts model | $\eta$ |
| dimension of the lower representation | $d$ |
| number of clusters | $q$ |

### Differential expression analysis and GO enrichment analysis

For differential expression analysis, we used the `FindAllMarkers` function in the R package Seurat (v4.3.0) to perform the Wilcoxon rank-sum test with gene expression at least a 0.25−fold difference (log-scale) between the target domain and others. We obtained the final DEGs for each domain using the adjusted p-value (Benjamin-Hochberg correction) with a cutoff of 0.05. For GO enrichment analysis, we first used the `select` function in the R package AnnotationDbi (v1.60.2) to transfer the gene symbol to entrezid. For the DEGs of each domain, we used the `enrichGO` function in the R package clusterProfiler (v4.6.2) to perform GO enrichment analysis for gene ontology over-representation test (one-sided version of Fisher’s exact test).

## Results

### STADIA enables more accurate correction of batch effects in the human dorsolateral prefrontal cortex dataset

To quantitatively evaluate the batch mixing and spatial clustering of STADIA, we first applied it to the human dorsolateral prefrontal cortex (DLPFC) dataset measured by the 10x Genomics Visium^38^. There are a total of 12 tissue slices from three independent neurotypical adult donors (**Fig. 2a**), each with two pairs of spatially adjacent replicates per adult (four slices per donor). All slices were manually labeled as layers 1 to 6 and white matter (WM) based on cytoarchitecture and genetic markers in the original publication, which will be used as ground truth to evaluate clustering accuracy.

**Fig. 2.**
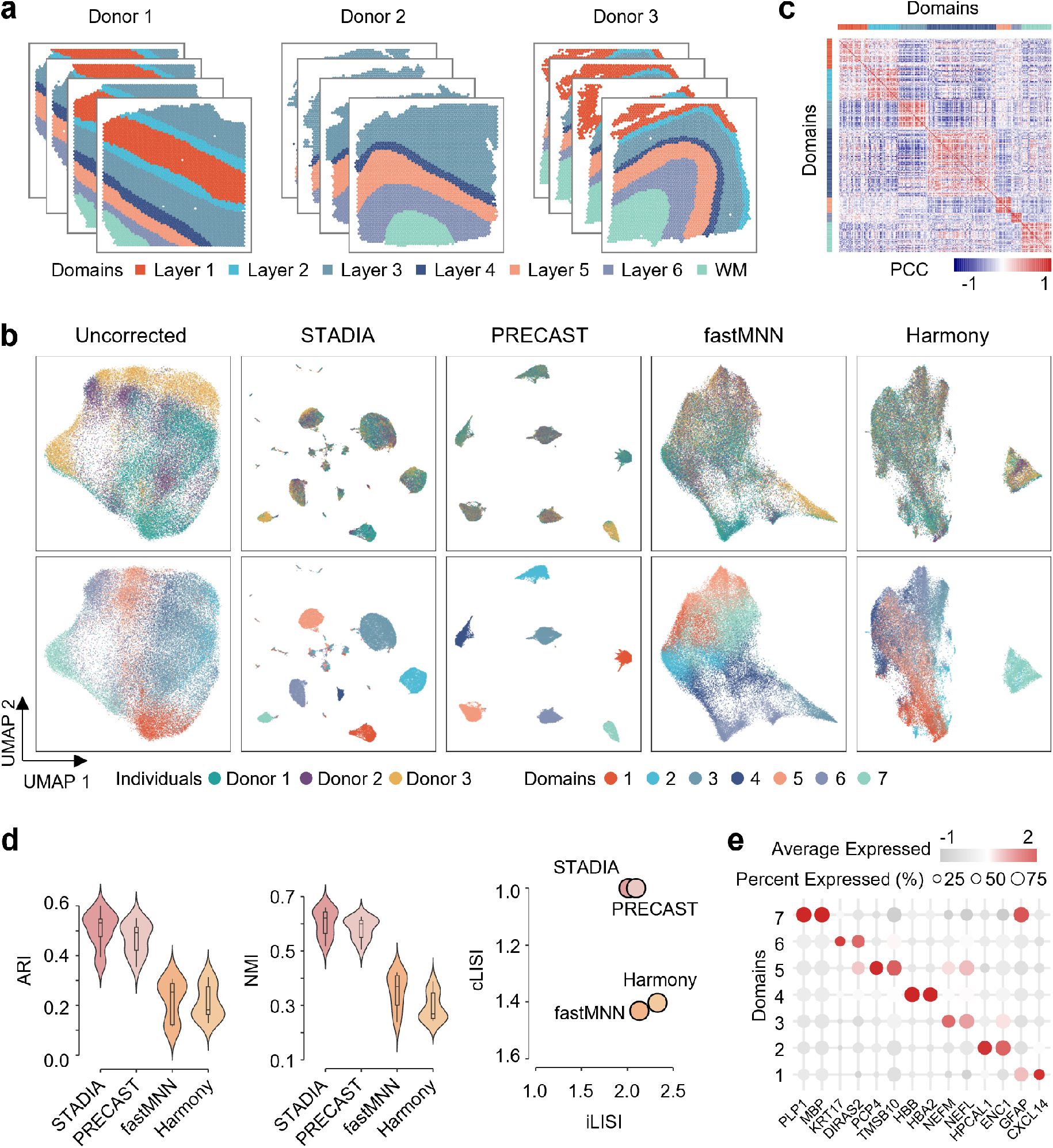
STADIA allows more accurate identification of layer structures and correction of batch effects in the human dorsolateral prefrontal cortex data set. **a**. Manual annotation of 12 slices from three donors based on cytoarchitecture labeled by layers 1 to 6 and white matter (WM). **b**. Uniform manifold approximation and projection (UMAP) visualization of the original data without correction, STADIA, PRECAST, fastMNN and Harmony, colored by the donor (top panel) and cluster assignments identified by the corresponding methods (bottom panel). **c**. Heatmap of Pearson’s correlation of the gene expressions between different spatial domains identified by STADIA. **d**. Violin plots of clustering accuracy in terms of ARI and UMI for the four methods (left and middle panels); scatter plot of mixing scores in terms of LISI, with batch mixing score along the x-axis and spatial domain mixing score along the y-axis (right panel). A point closer to the upper right corner indicates better performance. **e**. Dot plot of the top two marker genes for each spatial domain found by the Wilcoxon rank-sum test.

From the Uniform Manifold Approximation and Projection (UMAP) plot of the uncorrected raw data, there are substantial batch effects for the three different donors and negligible batch effects for the slices from the same donor (**Fig. 2b, upper left panel** and **Fig. S1a**). Therefore, we integrated all 12 slices using STADIA, PRECAST, and two commonly used batch effect correction strategies developed for scRNA-seq data, fastMNN and Harmony. From the embedded UMAP plots of these four methods, they all mixed the 12 slices well and had comparable Local Inverse Simpson’s Index (LISI) values (**Fig. 2b, top panel** and **Fig. 2d**, *x***-axis of the right panel**). Moreover, STADIA and PRECAST yielded more discriminative clusters than fastMNN and Harmony (**Fig. 2b, bottom panel** and **Fig. 2d**, *y***-axis of the right panel**), indicating that STADIA and PRECAST are adept at preserving the internal structure of the data while effectively distinguishing between different clusters. Next, comparing the spatial domains identified by the four methods with the ground truth, STADIA had the highest clustering accuracy in terms of Adjusted Rand Index (ARI) and Normalized Mutual Information (NMI), with a median ARI of 0.531 and a median NMI of 0.630, which were higher than PRECAST (median ARI = 0.492 and median NMI = 0.600), fastMNN (median ARI = 0.260 and median NMI = 0.367) and Harmony (median ARI = 0.161 and median NMI = 0.263) (**Fig. 2d, left and middle panel**). Furthermore, spatial visualization of all slices demonstrated that STADIA exhibited more consistent spatial domains compared to alternative methods. Additionally, STADIA showcased smoother layer boundaries (**Fig. S1b**), aligning more consistently with the coherence and structural integrity of the cortical hierarchy. We also calculated Pearson correlations of all clusters to verify the plausibility of the spatial domains identified by STADIA (**Fig. 2c**). Finally, by comparing each layer with all other layers using the Wilcoxon rank-sum test, we identified marker genes that were differentially expressed in each layer, including previously published markers: *HPCAL1* and *ENC1* for layer 2, *PCP4* for layer 5, *KRT17* for layer 6, and *MBP* for WM^38^ (**Fig. 2e** and **Fig. S1c**).

### STADIA enables the horizontal integration of two adjacent sagittal mouse brain slices while preserving slice-specific biological variation

In addition to multiple duplicate slices of a single section, there are many repetitions of horizontally adjacent slices in an experiment, such as data from sagittal mouse brain slices sequenced with the 10x Visium platform. This dataset consists of an anterior sagittal slice and a posterior sagittal slice. We first displayed the hematoxylin & eosin (H&E) stained images corresponding to sagittal mouse brain slices and the tissue structure of the Allen Mouse Brain Atlas (**Fig. 3a**) and zoomed in on the known hippocampus.

**Fig. 3.**
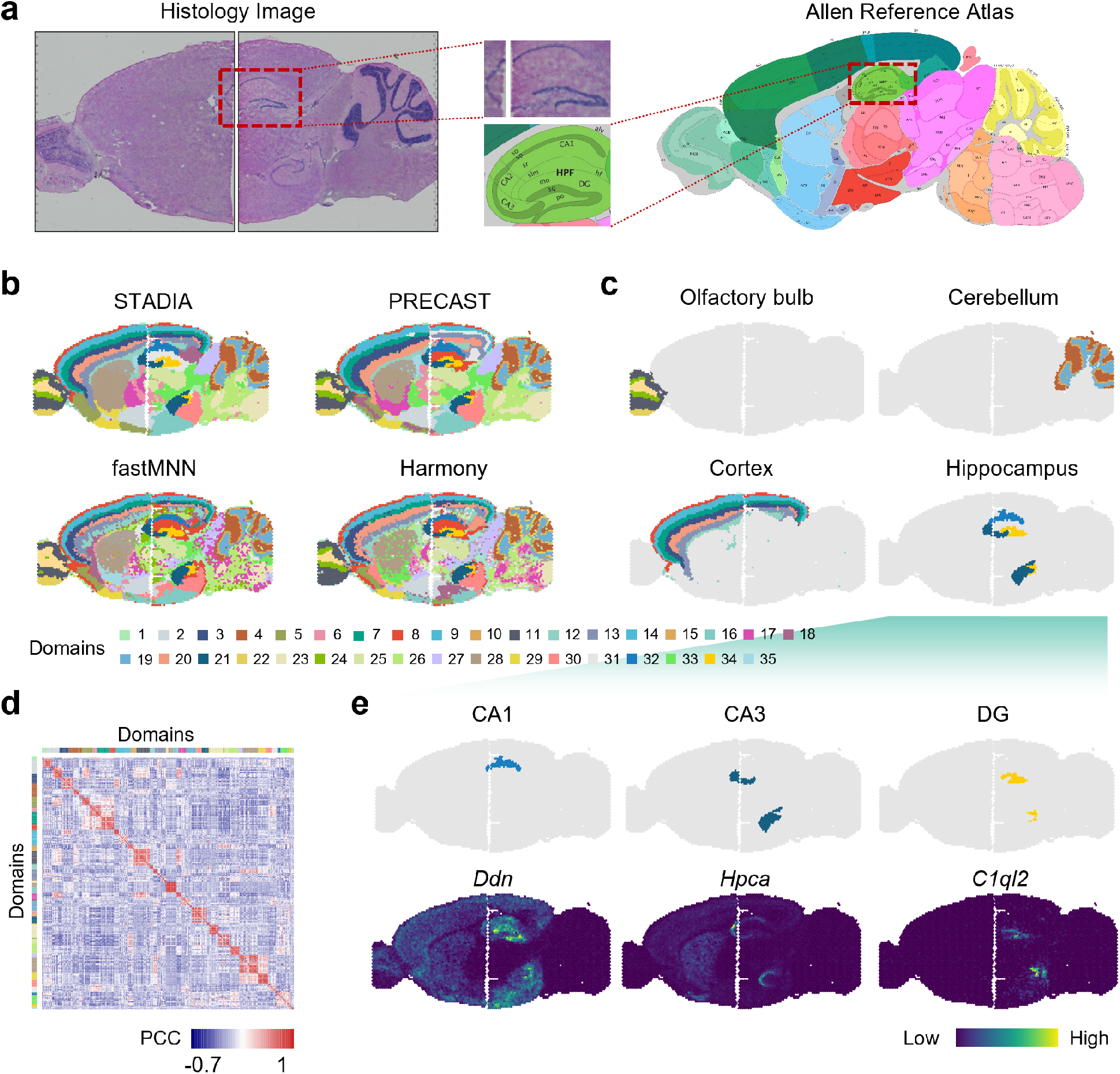
STADIA enables the horizontal integration of two adjacent mouse brain slices while preserving slice-specific biological variation. **a**. The Hematoxylin- and Eosin (H&E) images of two slices and the corresponding tissue structures obtained from the Allen Mouse Brain Atlas. **b**. Horizontal alignment of spatial domains of adjacent mouse brain slices, identified by STADIA, PRECAST, fastMNN and Harmony, respectively. **c**. Spatial visualization of slice-specific tissue structures olfactory bulb and cerebellum (top panel), slice-sharing tissue structures cortex and hippocampus (bottom panel), learned by STADIA. **d**. Heatmap of Pearson’s correlation of spatial domains identified by STADIA. **e**. Visualization of hippocampal subregions learned by STADIA and their corresponding top one marker genes found by the Wilcoxon rank-sum test.

From the full view, STADIA identified commonly known layer structures and horizontally aligned cluster assignments well across two slices (**Fig. 3b, upper left panel**), such as the shared organizational structures of the cerebral cortex layer (domains 3, 7, 8, 9, 13, 20) and the hippocampus (domains 21, 32, 34) (**Fig. 3c, bottom panel**). This was further validated by the expression of cluster-specific markers *Ddn, Hpca* and *C1ql2* of the hippocampal subregions cornu ammonis 1 (CA1), cornu ammonis 3 (CA3) and dentate gyrus (DG), respectively (**Fig. 3e**). While identifying the shared tissue structures, STADIA also preserved the slice-specific biological variations, such as the slice-specific tissue structures *olfactory bulb* and *cerebellum* (**Fig. 3c, top panel**). In comparison, the spatial partitioning of fastMNN and Harmony showed considerable noise, no clear layer boundaries, and PRECAST did not well align domains between the two slices, such as the cortex (**Fig. 3b**). To further evaluate STADIA, we calculated Pearson correlations between all domains (**Fig. 3d**), demonstrating a significantly higher within-group similarity compared to between-group similarity.

### STADIA enables the detection of person-specific cancer domains verified by cancer-associated markers

Despite having the same type of cancer, patients show very very different symptoms. In this section, we studied the human cutaneous squamous cell carcinoma (cSCC) dataset^39^ processed according to the ST protocol^40^. It consisted of 12 slices from different parts of four patients, with three cryosections per patient. Specifically, these samples were taken from the left forearm, left vertex scalp, right forearm, and right tragus of each individual (**Fig. 4a**).

**Fig. 4.**
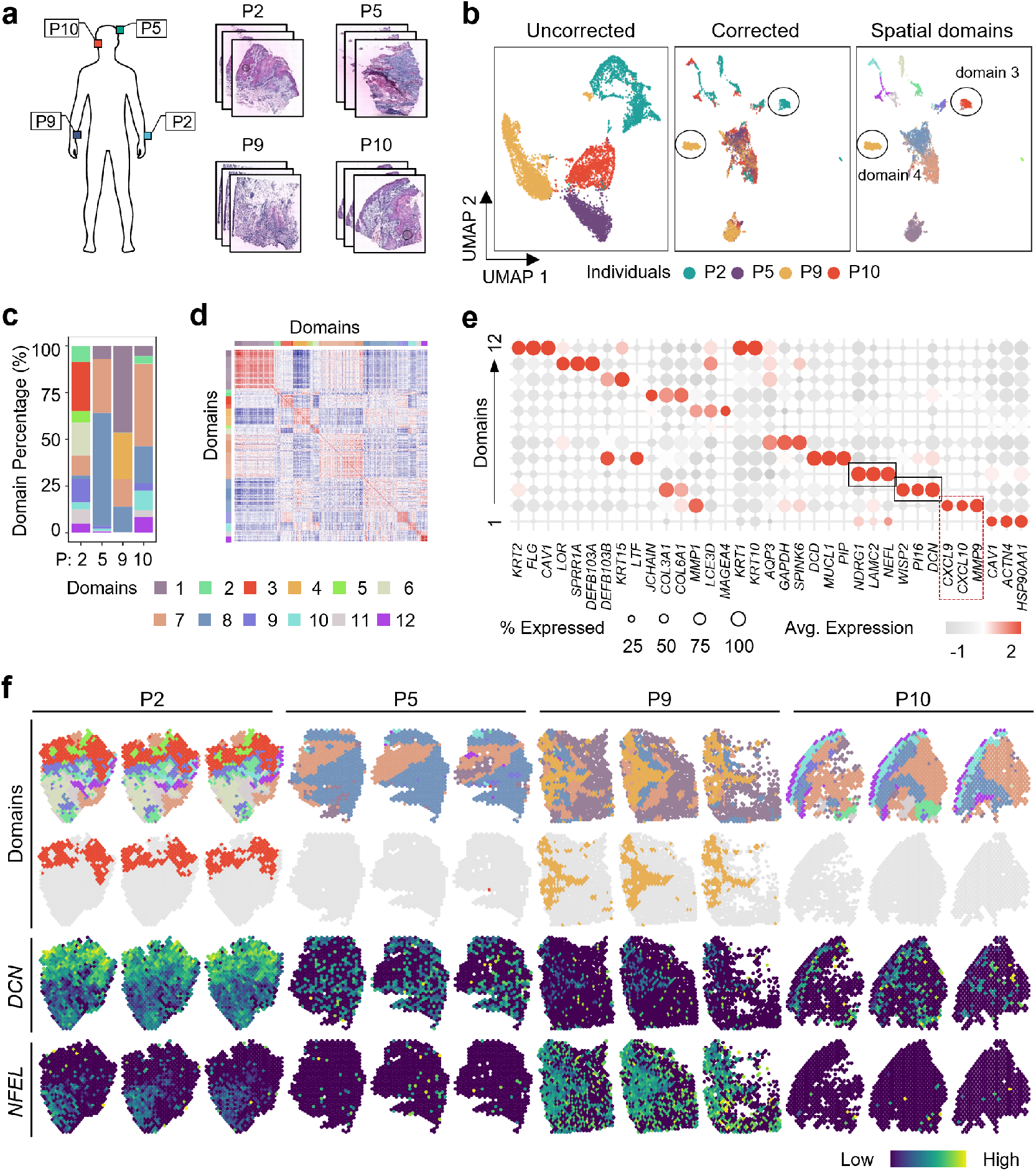
STADIA enables the detection of person-specific cancer domains verified by cancer-associated markers in the cSCC dataset. **a**. Hematoxylin and Eosin (H&E) images of 12 cSCC slices from different parts of four patients. The top-left three slices were from the left forearm of P2, the top-right three slices were from the left vertex scalp of P5, the bottom-left three slices were from the right forearm of P9, and the bottom-right three slices were from the right tragus of P10. **b**. UMAP plots of the original data without correction colored by patients (left panel), and embeddings for STADIA colored by patients (middle panel) and cluster assignments (right panel). **c**. Percentage distribution of spatial domains for the four patients, colored as in the left panel of (b). **d**. Heatmap of Pearson’s correlation of the gene expressions between different spatial domains identified by STADIA. **e** Dot plot of the top three markers for each spatial domain identified by STADIA. **f**. Visualization of spatial domains identified by STADIA in a spatial context (top panel), with domains 3 and 4 highlighted (second panel) and spatial visualization of the corresponding marker genes *CND* for domain 3 (overexpressed in patient P2) and *NEEL* for domain 4 (overexpressed in P9) (right panel).

From the UMAP plot of the uncorrected raw data, there was little overlap in these four patients, but the batch effect between the different slices of each patient is not significant (**Fig. 4b, left panel** and **Fig. S2a**). Based on prior knowledge of the cancers, the lack of overlap may be due to tumor heterogeneity and some degree of batch effects. After correction by STADIA, the embeddings were well mixed, while there were some isolated domains for the P2 and P9 patients, such as domain 3 for P2 and domain 4 for P9 (**Fig. 4b, middle and right panels**).

Although all slices were cSCC samples, the cell-type composition of these four patients varied considerably (**Fig. 4c**), suggesting that these patients may be at different stages of disease progression or have different cancer subtypes. To further explore the correlation of these domains, we calculated the Pearson correlations across all domains (**Fig. 4d**). The samples from patient P9 mainly consisted of domain 1 and domain 4, which were negatively correlated (**Fig. 4d**). The most significantly differentially expressed gene in domain 1 was *HSP90AA1* (**Fig. 4e**), which is associated with disease progression and potential clinical targets for SCC patients^41^. In particular, domain 4 was uniquely present in P9, which was confirmed by the marker gene *NEFL* (**Fig. 4e and Fig. 4f**). Similarly, domain 3 appeared only in P2 with high expression of the gene *DCN* (**Fig. 4e** and **Fig. 4f**) and expressed markers of fibroblasts such as *PI16* and *WISP2*.

Previous studies have shown that the expression of both *DCN* and *NEFL* correlates with the invasive ability of cancer cells and that high levels of *DCN* and *NEFL* decrease the invasive ability of cancer cells^42,43^. In contrast, highly expressed marker genes for domain 2 were therapeutic targets for related human malignancies, such as *MMP9*^44,45^, *CXCL10*^46^, *CXCL9*^47^, suggesting that domain 2 is a relatively severe tumor region (**Fig. 4e**). Furthermore, from the stacked bar plot of cell-type composition, we can directly conclude that domain 2 was predominantly found in P2 and P10 patients (**Fig. 4c**). To refine our analysis, we visualized the expression patterns of the top three markers within domain 2 in a spatial context. The results showed that *MMP9* was highly expressed in domain 2 of both P2 and P10, while *CXCL10* and *CXCL9* exhibited elevated expression levels exclusively in domain 2 of P2 (**Fig. S2b**). This observation was further confirmed by the violin plots of the expression of these three genes (**Fig. S2c**). Such distinctions in expression profiles may suggest the presence of distinct cellular subtypes or functional subtypes within domain 2 in these two patients, which could be associated with different characteristics or clinical factors of the tumor.

### STADIA learns the common biological variations among eight mouse liver tissue sections

We further applied STADIA to the mouse liver dataset^48^, which consists of eight liver tissue slices from three adult female wild mice profiled by the ST protocol. Six slices were from the caudate lobe portion of mouse 1 (CN73) and mouse 2 (CN65) (three slices per mouse), and two were from the right lobe portion of mouse 3 (CN16) (**Fig. 5a**).

**Fig. 5.**
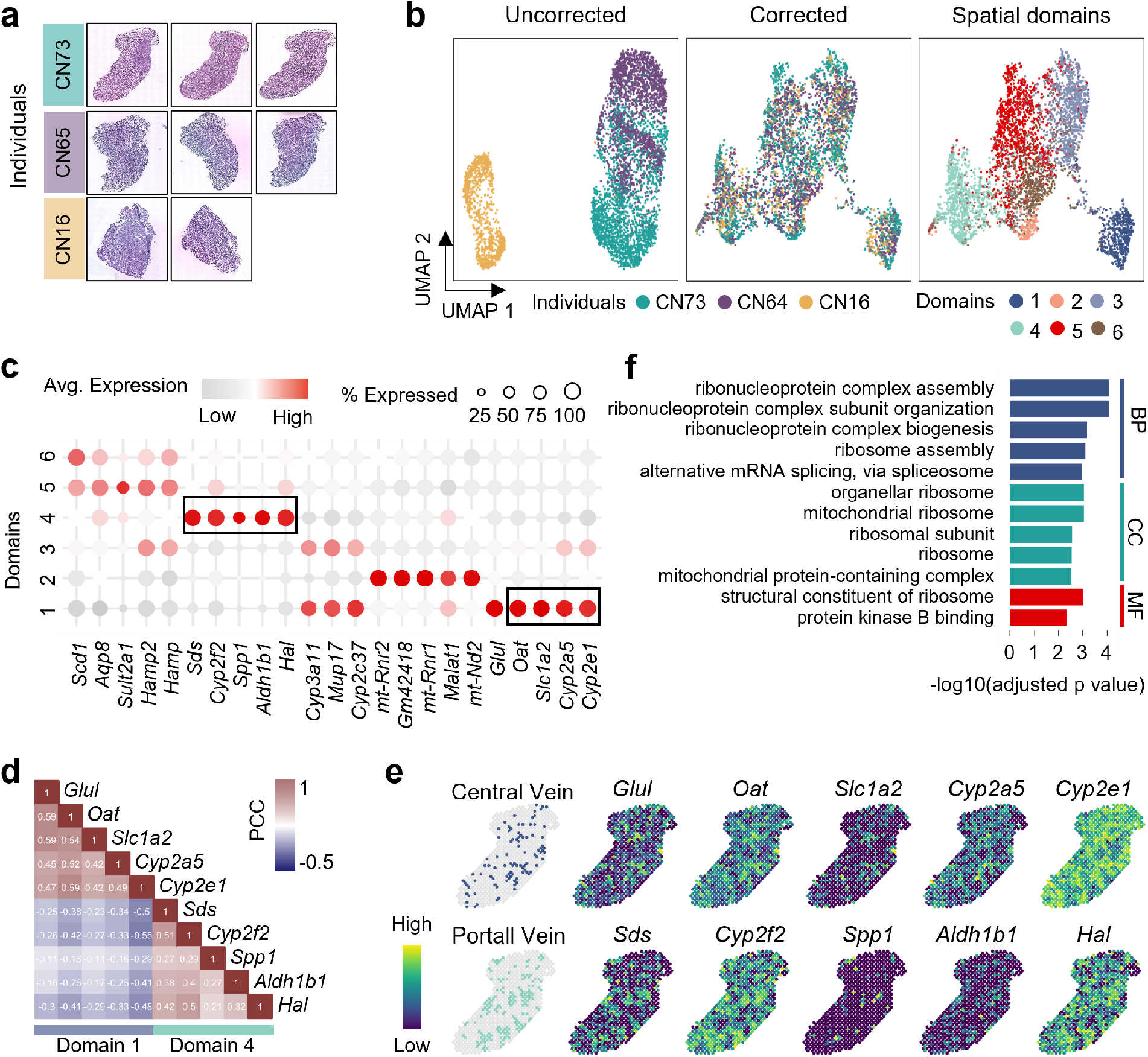
STADIA learns the common biological variations among eight mouse liver tissue sections. **a**. Hematoxylin and Eosin (H&E) images for all eight mouse liver tissue sections. Slices from the first two samples (CN73, CN65) are from parts of the caudate lobe, and slices from the last sample (CN16) are from parts of the right lobe. **b**. UMAP plots of the original data without correction, colored by mouse (left panel), UMAP plots of embeddings for STADIA, colored by mouse (middle panel), and cluster assignment (right panel). **c**. Dot plot of top five found by the Wilcoxon rank-sum test for each spatial domain identified by STADIA. **d**. Heatmap of Pearson’s correlation of the marker gene expression profiles for domains 1 and 4. **e**. Spatial distribution of the marker genes for spatial domains 1 (central vein) and 4 (portal vein) as listed in (c) and (d). **f**. Bar chart of the GO enrichment analysis results for spatial domain 2. The enrichment score is calculated as the -log10(adjusted p-value).

From the UMAP plot of the uncorrected raw data, there were significant differences between the three mice, with individual CN73 being closer to CN65 than CN16, which may because both CN73 and CN65 were sampled from the caudate lobe (**Fig. 5b, left panel**). Batch effects across slices from the same individual were negligible (**Fig. S3a**). In addition, STADIA mixed all eight slices well and separated the six domains on the embedded UMAP plot (**Fig. 5b, middle and right panels**). Using the Wilcoxon rank-sum test, we found the domain-specific marker genes and plotted the top five for each domain in a dot plot (**Fig. 5c**). The top five marker genes for domain 1 and domain 4 included previously published zoned hepatocyte genes *Glul, Cyp2e1, Sds*, and *Hal*^49^, suggesting that domain 1 and domain 4 are central vein (CV) and portal vein (PV), respectively. To further confirm our findings, we calculated Pearson correlations of marker genes for domain 1 and domain 4 (**Fig. 5d** and **Fig. S3c**) and visualized their expression in a spatial context (**Fig. 5e**), which indicated that the marker gene expression of these two organizational structures was negatively correlated.

We further explored the biological significance of differentially expressed genes (DEGs) by Gene Ontology (GO) enrichment analysis in terms of biological process (BP), cellular component (CC) and molecular function (MF). Among them, focusing on up-regulated DEGs for domain 2, the enriched biological processes were mainly related to ribonucleoprotein complex assembly and ribonucleoprotein complex biogenesis. In terms of cellular components, DEGs were enriched in ribosomes of organelles and ribosomes. In terms of molecular function, only the terms for structural constituent of ribosome and protein kinase B binding were enriched (**Fig. 5f**). DEGs for other domains were mainly enriched in different metabolic pathways (**Fig. S3d**).

### STADIA allows identification of common hippocampal tissue structures while preserving slice-specific biological variation by integrating two hippocampal slices from Slide-seqV2

To demonstrate the scalability of STADIA to different spatial resolutions, we applied STADIA to a mouse hippocampus dataset profiled by Slide-seqV2^50^, which can profile spatial expression at near-cellular resolution (10 µm). This dataset consists of two slices with batch effects present (**Fig. S4a, upper left panel**).

From the Allen Reference Atlas, we can see that the hippocampal tissue consists of the cornu ammonis (CA) with subregions CA1, CA2 and CA3, and the dentate gyrus (DG) (**Fig. 6a**). As expected, STADIA removed batch effects of the two slices (**Fig. S4a**) and successfully characterized the hippocampal structures for both slices with clear domain boundaries (**Fig. 6b**). In addition, the domain-specific marker genes were found by the Wilcoxon rank-sum test, and five of all common domains together with their most significant marker genes were plotted in a spatial context (**Fig. 6c**). For example, the expression of *C1ql2* showed a clear arrowhead shape that was highly expressed in the hippocampal subregion DG and the expression of *Ddn* was highly expressed around the structure of the hippocampus corresponding to domain 5.

**Fig. 6.**
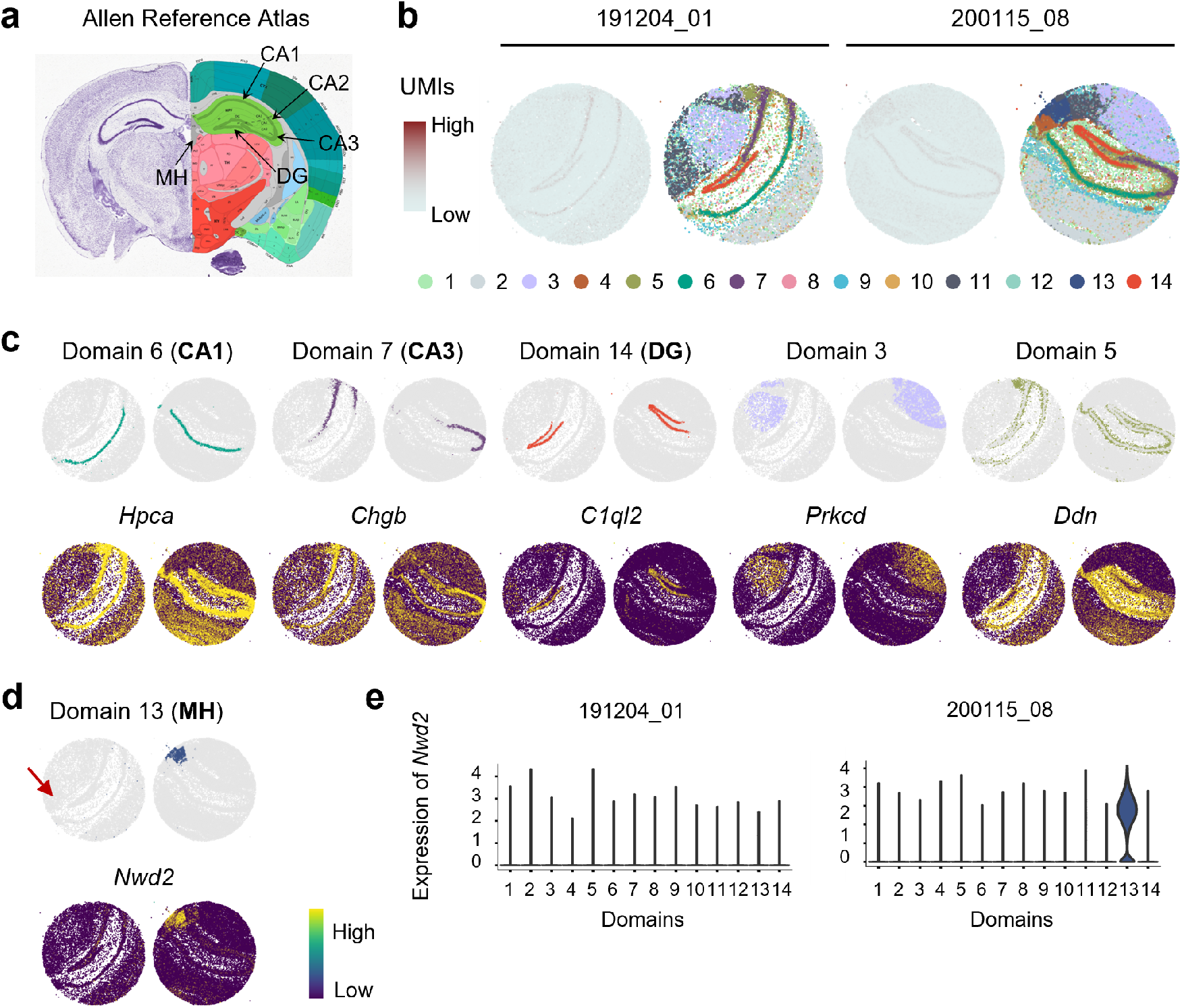
STADIA enables the identification of common hippocampal tissue structures while preserving slice-specific variation by integrating two hippocampal slices from Slide-seqV2. **a**. The reference tissue structures from the Allen Mouse Brain Atlas. **b**. Spatial visualization for two slices, colored by the number of UMIs and spatial domains identified by STADIA. **c**. Spatial visualization of domains 6, 7, 14, 3 and 5 identified by STADIA with their top marker genes found by the Wilcoxon rank-sum test. **d**. The slice-specific domain 13 identified by STADIA for the second slice, marked by the gene *Nwd2*. **e**. The violin plots for the expression of the gene *Nwd2* of two slices. **f**. Histogram of the log2-transformed number of spots and UMIs for two slices, with vertical lines corresponding to the gene *Nwd2*.

Furthermore, a slice-specific domain (domain 13) was identified by STADIA for the second slice (puck_200115_08), which was characterized by the marker gene *Nwd2* (**Fig. 6d**). We found that *Nwd2* was highly expressed only in domain 13 of the second slice (**Fig. 6e**). This underscores STADIA’s ability to maintain the distinct biological variations of individual slices while uncovering shared biological properties across multiple slices.

## Discussion

Due to the limitations of existing technologies, a single experiment can only capture limited biological signals in a small region. With the rapid accumulation of ST data, we developed a hierarchical hidden Markov random field model STADIA to align spots from multiple ST slices with batch-effect correction. The observed expression profiles and the batch-corrected embeddings are linked by a Bayesian factor regression model with L/S adjustment. The batch-corrected embeddings and the cluster assignments are linked by a GMM model, with the cluster assignments further modeled by the Potts model to ensure local smoothing. Through extensive experiments, STADIA can successfully mix multiple ST slices without overcorrection and preserve slice-specific biological variations.

The main limitation is that the linear dimensionality reduction we used results in more data loss than a nonlinear method. Future work is expected to employ nonlinear methods such as an autoencoder, which links the observed expression to hidden embeddings, and Markov random field model-driven clustering methods for further spatially aware embedded clustering.

Our current work is limited to transcriptome analysis. In the future, we plan to jointly analyze muti-omics data across multiple slices, which can provide insights into the regulation of the entire process, spanning from gene to protein expression. To address this issue, we need to consider the batch effects across different slices as well as the regulatory mechanisms of various omics, which poses a greater challenge.

## Data Availability

All datasets analyzed in this study are available through websites reported in the original publications. Specifically, the Human dorsolateral prefrontal cortex (DLPFC) data can be accessed in the spatialLIBD package, which is available at http://research.libd.org/spatialLIBD/. The mouse sagittal posterior and anterior brain data can be accessed in official 10x genomics support https://support.10xgenomics.com/spatial-gene-expression/datasets/1.0.0/V1_Mouse_Brain_Sagittal_Anterior, https://support.10xgenomics.com/spatial-gene-expression/datasets/1.0.0/V1_Mouse_Brain_Sagittal_Posterior. The cutaneous squamous cell carcinoma (cSCC) data can be accessed at the Gene Expression Omnibus (GEO) under the accession code GSE144240.

The mouse liver data are available in the DOI-minting Zenodo repository 5595907. The mouse hippocampus dataset can be accessed at https://singlecell.broadinstitute.org/single_cell/study/SCP815. The image of Allen Mouse Brain Atlas can be accessed at http://atlas.brain-map.org/atlas?atlas=2&plate=100883818.

## Code Availability

The STADIA algorithm is implemented using R software and is packaged as an R package *stadia*, which is available at https://github.com/zhanglabtools/STADIA.

## Acknowledgements

This work has been supported by the National Key Research and Development Program of China [No. 2021YFA1302500 to S.Z.], the National Natural Science Foundation of China [No. 12126605], the Key-Area Research and Development of Guangdong Province [No. 2020B1111190001], and the CAS Project for Young Scientists in Basic Research [No. YSBR-034 to S.Z.], and the China Postdoctoral Science Foundation [No. 2022M723328 to Y.L.].

## Author contributions

S.Z. conceived and supervised the project. Y.L. developed and implemented the STADIA algorithm. Y.L. and S.Z. validated the methods and wrote the manuscript. All authors read and approved the final manuscript.

## Competing interests

The authors declare no competing interests.

## Notes

### Competing Interest Statement

The authors have declared no competing interest.

## References

1. Zhao, E. et al. Spatial transcriptomics at subspot resolution with BayesSpace. Nature Biotechnology 39, 1375–1384 (2021).

2. Dong, K. & Zhang, S. Deciphering spatial domains from spatially resolved transcriptomics with an adaptive graph attention auto-encoder. Nature Communications 13, 1–12 (2022).

3. Hu, J. et al. SpaGCN: Integrating gene expression, spatial location and histology to identify spatial domains and spatially variable genes by graph convolutional network. Nature Methods 18, 1342–1351 (2021).

4. Fu, H. et al. Unsupervised spatially embedded deep representation of spatial transcriptomics. bioRxiv, 2021–06 (2021).

5. Liu, W. et al. Joint dimension reduction and clustering analysis of single-cell RNA-seq and spatial transcriptomics data. Nucleic Acids Research 50, e72–e72 (2022).

6. Edsgärd, D., Johnsson, P. & Sandberg, R. Identification of spatial expression trends in single-cell gene expression data. Nature Methods 15, 339–342 (2018).

7. Svensson, V., Teichmann, S. A. & Stegle, O. SpatialDE: identification of spatially variable genes. Nature Methods 15, 343–346 (2018).

8. Sun, S., Zhu, J. & Zhou, X. Statistical analysis of spatial expression patterns for spatially resolved transcriptomic studies. Nature Methods 17, 193–200 (2020).

9. Zhu, J., Sun, S. & Zhou, X. SPARK-X: non-parametric modeling enables scalable and robust detection of spatial expression patterns for large spatial transcriptomic studies. Genome Biology 22, 1–25 (2021).

10. Andersson, A. & Lundeberg, J. sepal: identifying transcript profiles with spatial patterns by diffusion-based modeling. Bioinformatics 37, 2644–2650 (2021).

11. Zhang, C., Dong, K., Aihara, K., Chen, L. & Zhang, S. STAMarker: determining spatial domain-specific variable genes with saliency maps in deep learning. Nucleic Acids Research, gkad801 (2023).

12. Elosua-Bayes, M., Nieto, P., Mereu, E., Gut, I. & Heyn, H. SPOTlight: seeded NMF regression to deconvolute spatial transcriptomics spots with single-cell transcriptomes. Nucleic Acids Research 49, e50–e50 (2021).

13. Shan, X., Chen, J., Dong, K., Zhou, W. & Zhang, S. Deciphering the spatial modular patterns of tissues by integrating spatial and single-cell transcriptomic data. Journal of Computational Biology 29, 650–663 (2022).

14. Ma, Y. & Zhou, X. Spatially informed cell-type deconvolution for spatial transcriptomics. Nature Biotechnology 40, 1349–1359 (2022).

15. Lu, Y., Chen, Q. & An, L. SPADE: Spatial Deconvolution for Domain Specific Cell-type Estimation. bioRxiv, 2023–04 (2023).

16. Johnson, W. E., Li, C. & Rabinovic, A. Adjusting batch effects in microarray expression data using empirical Bayes methods. Biostatistics 8, 118–127 (2007).

17. Korsunsky, I. et al. Fast, sensitive and accurate integration of single-cell data with Harmony. Nature Methods 16, 1289–1296 (2019).

18. Haghverdi, L., Lun, A. T., Morgan, M. D. & Marioni, J. C. Batch effects in single-cell RNA-sequencing data are corrected by matching mutual nearest neighbors. Nature Biotechnology 36, 421–427 (2018).

19. Lun, A. Further MNN algorithm development. https://MarioniLab.github.io/FurtherMNN2018/theory/description.html. (2019).

20. Hie, B., Bryson, B. & Berger, B. Efficient integration of heterogeneous single-cell transcriptomes using Scanorama. Nature Biotechnology 37, 685–691 (2019).

21. Hardoon, D. R., Szedmak, S. & Shawe-Taylor, J. Canonical correlation analysis: An overview with application to learning methods. Neural Computation 16, 2639–2664 (2004).

22. Stuart, T. et al. Comprehensive integration of single-cell data. Cell 177, 1888–1902 (2019).

23. Zeira, R., Land, M., Strzalkowski, A. & Raphael, B. J. Alignment and integration of spatial transcriptomics data. Nature Methods 19, 567–575 (2022).

24. Zhou, X., Dong, K. & Zhang, S. Integrating spatial transcriptomics data across different conditions, technologies and developmental stages. Nature Computational Science, 1–13 (2023).

25. Long, Y. et al. Spatially informed clustering, integration, and deconvolution of spatial transcriptomics with GraphST. Nature Communications 14, 1155 (2023).

26. Luo, X. & Wei, Y. Batch effects correction with unknown subtypes. Journal of the American Statistical Association 114, 581–594 (2019).

27. Avalos-Pacheco, A., Rossell, D. & Savage, R. S. Heterogeneous large datasets integration using Bayesian factor regression. Bayesian Analysis 17, 33–66 (2022).

28. Liu, W. et al. Probabilistic embedding, clustering, and alignment for integrating spatial transcriptomics data with PRECAST. Nature Communications 14, 296 (2023).

29. Fraley, C. & Raftery, A. E. Model-based clustering, discriminant analysis, and density estimation. Journal of the American statistical Association 97, 611–631 (2002).

30. McLachlan, G. J., Lee, S. X. & Rathnayake, S. I. Finite mixture models. Annual review of statistics and its application 6, 355–378 (2019).

31. Graner, F. & Glazier, J. A. Simulation of biological cell sorting using a two-dimensional extended Potts model. Physical Review Letters 69, 2013 (1992).

32. Hao, Y. et al. Integrated analysis of multimodal single-cell data. Cell 184, 3573–3587 (2021).

33. Liu, C. & Rubin, D. B. ML estimation of the t distribution using EM and its extensions, ECM and ECME. Statistica Sinica 5, 19–39 (1995).

34. Schuurman, N., Grasman, R. & Hamaker, E. A comparison of inverse-wishart prior specifications for covariance matrices in multilevel autoregressive models. Multivariate Behavioral Research 51, 185–206 (2016).

35. Gottardo, R., Besag, J., Stephens, M. & Murua, A. Probabilistic segmentation and intensity estimation for microarray images. Biostatistics 7, 85–99 (2006).

36. Johnson, V. E. & Rossell, D. On the use of non-local prior densities in Bayesian hypothesis tests. Journal of the Royal Statistical Society: Series B (Statistical Methodology) 72, 143–170 (2010).

37. Johnson, V. E. & Rossell, D. Bayesian model selection in high-dimensional settings. Journal of the American Statistical Association 107, 649–660 (2012).

38. Maynard, K. R. et al. Transcriptome-scale spatial gene expression in the human dorsolateral prefrontal cortex. Nature Neuroscience 24, 425–436 (2021).

39. Ji, A. L. et al. Multimodal analysis of composition and spatial architecture in human squamous cell carcinoma. Cell 182, 497–514 (2020).

40. Ståhl, P. L. et al. Visualization and analysis of gene expression in tissue sections by spatial transcriptomics. Science 353, 78–82 (2016).

41. Fan, G., Tu, Y., Wu, N. & Xiao, H. The expression profiles and prognostic values of HSPs family members in Head and neck cancer. Cancer cell international 20, 1–12 (2020).

42. Hu, X. et al. Decorin-mediated suppression of tumorigenesis, invasion, and metastasis in inflammatory breast cancer. Communications Biology 4, 72 (2021).

43. Huang, Z. et al. The role of NEFL in cell growth and invasion in head and neck squamous cell carcinoma cell lines. Journal of Oral Pathology & Medicine 43, 191–198 (2014).

44. Augoff, K., Hryniewicz-Jankowska, A., Tabola, R. & Stach, K. MMP9: a tough target for targeted therapy for cancer. Cancers 14, 1847 (2022).

45. Tufaro, A. P. et al. Molecular markers in cutaneous squamous cell carcinoma. International Journal of Surgical Oncology 2011 (2011).

46. Liu, M., Guo, S. & Stiles, J. K. The emerging role of CXCL10 in cancer. Oncology Letters 2, 583–589 (2011).

47. Ding, Q. et al. CXCL9: evidence and contradictions for its role in tumor progression. Cancer Medicine 5, 3246–3259 (2016).

48. Hildebrandt, F. et al. Spatial Transcriptomics to define transcriptional patterns of zonation and structural components in the mouse liver. Nature Communications 12, 7046 (2021).

49. Guilliams, M. et al. Spatial proteogenomics reveals distinct and evolutionarily conserved hepatic macrophage niches. Cell 185, 379–396 (2022).

50. Stickels, R. R. et al. Highly sensitive spatial transcriptomics at near-cellular resolution with Slide-seqV2. Nature Biotechnology 39, 313–319 (2021).

