## Supplemental Text and Figures for "Statistical batch-aware embedded integration, dimension reduction and alignment for spatial transcriptomics"

### Supplementary Notes

#### Marginal student-t distribution [Liu & Rubin[1]]

Multivariate t-distribution $t_d(\nu, \boldsymbol{\mu}, \boldsymbol{\Sigma})$ is a multivariate probability distribution with density

$$f_t(\boldsymbol{x}; \nu, \boldsymbol{\mu}, \boldsymbol{\Sigma}) = \frac{\Gamma\left(\frac{\nu+d}{2}\right)}{\Gamma\left(\frac{\nu}{2}\right)(\nu\pi)^{\frac{d}{2}} |\boldsymbol{\Sigma}|^{\frac{1}{2}}} \left[1 + \frac{1}{\nu}(\boldsymbol{x} - \boldsymbol{\mu})^\top \boldsymbol{\Sigma}^{-1}(\boldsymbol{x} - \boldsymbol{\mu})\right]^{-\frac{\nu+d}{2}}.$$

**Lemma 0.1.** *When given the weight $\tau$, $\boldsymbol{x}$ has the multivariate normal distribution*

$$\boldsymbol{x} \mid \tau \sim \mathcal{N}_p(\boldsymbol{\mu}, \tau^{-1}\boldsymbol{\Sigma}),$$

*and $\tau\nu$ is $\chi^2_\nu$, that is, the weight $\tau$ is Gamma distribution given $\nu$*

$$\tau \mid \nu \sim \textit{Gamma}(\nu/2, \nu/2),$$

*then the marginal distribution of $\boldsymbol{x}$ by integrating out $\tau$ is multivariate student-t distribution $t_p(\nu, \boldsymbol{\mu}, \boldsymbol{\Sigma})$.*

#### Motivation of L/S adjustment

The Location-and-Scale (L/S) adjustment is the cornerstone of the ComBat algorithm applied to microarray data for batch effect correction, which works by estimating batch-specific location and scale parameters for each gene or feature across various batches. Similarly, STADIA adopts a similar strategy to harmonize gene expression across different batches within spatial transcriptomics (ST) datasets. Here, we outline the rationale behind the L/S adjustment from two aspects.

1. Experimental aspect: Systematic differences between batches, such as variations in instrument calibration and batches of reagents, can lead to an overall shift in the measurement results. Additionally, factors like experimental platforms and instrument sensitivity may cause changes in data proportions, even when measuring the same feature in the same samples. The L/S adjustment addresses these challenges by employing additive and multiplicative corrections. It adds a batch-specific bias and scaling factor to each variable, effectively correcting for shifts and proportional changes induced by the batch effects.

2. Data distribution aspect: After normalization, data generally follows a normal distribution, uniquely determined by its mean and covariance matrix. The mean represents the central tendency of the distribution, while the covariance matrix measures the variation of the randomness. To harmonize the gene expressions across different batches, which are assumed to be normally distributed, adjusting the means and covariance matrices is sufficient.

#### Overall framework of Bayesian inference

First, all parameters and all latent or missing variables are combined into $\theta = \{\mathbf{L}, \mathbf{T}, \boldsymbol{\Lambda}, \boldsymbol{\gamma}, \boldsymbol{\mu}, \boldsymbol{\omega}, \mathbf{p}\}$ and $\mathbf{Z} = \{\boldsymbol{f}, \boldsymbol{c}, \boldsymbol{s}\}$ respectively. The Expectation-Maximization (EM) algorithm is then used to obtain maximum a posteriori (MAP) estimates for all parameters. By standard algebraic calculations with Jensen's Inequality, a tight lower bound for the log posterior is given by

$$\log\left[\mathrm{P}(\theta \,|\, \mathbf{X})\right] \geqslant \mathbb{E}_{z|x,\theta^{(t)}} \log\left[\mathrm{P}(\theta \,|\, \mathbf{X}, \mathbf{Z})\right] + C \overset{\text{def}}{=} Q(\theta \,|\, \theta^{(t)}) + C, \tag{0.1}$$

where $\mathbb{E}_{z|x,\theta^{(t)}}$ denotes the expectation of the latent variable $\mathbf{Z}$ conditional on the observation $\mathbf{X}$ and the current values of parameters $\theta^{(t)}$, $C$ is a constant with respect to $\theta$, and all of the following $C$'s are constants whose values vary from place to place. The EM algorithm then iteratively solves the following optimization problem

$$\theta^{(t+1)} = \arg\max_{\theta} Q(\theta \,|\, \theta^{(t)}).$$

In the expectation step, the core objective is to obtain an explicit notation for $Q(\theta \,|\, \theta^{(t)})$, by computing the two conditional distributions $\mathrm{P}(\theta \,|\, \mathbf{X}, \mathbf{Z})$ and $\mathrm{P}(\mathbf{Z} \,|\, \mathbf{X}, \theta^{(t)})$. From the conditional independence of $\{\boldsymbol{f}, \boldsymbol{c}, \boldsymbol{s}\}$ given others, we have

$$\begin{aligned}\mathrm{P}(\mathbf{Z} \,|\, \mathbf{X}, \theta^{(t)}) &= \prod_{b,i} \mathrm{P}(\boldsymbol{f}_{bi} \,|\, c_{bi}, \boldsymbol{x}_{bi}, \theta^{(t)}) \mathrm{P}(\boldsymbol{c} \,|\, \mathbf{X}, \theta^{(t)}) \mathrm{P}(\boldsymbol{s} \,|\, \theta^{(t)}) \\ &\approx \underbrace{\prod_{b,i} \mathrm{P}(\boldsymbol{f}_{bi} \,|\, c_{bi}, \boldsymbol{x}_{bi}, \theta^{(t)})}_{P_{11}} \underbrace{\prod_{b,i} \mathrm{P}\left(c_{bi} \;\middle|\; \hat{\boldsymbol{c}}_{\mathsf{N}_{bi}}, \boldsymbol{x}_{bi}, \theta^{(t)}\right)}_{\widetilde{P}_{12}} \underbrace{\mathrm{P}(\boldsymbol{s} \,|\, \theta^{(t)})}_{P_2},\end{aligned} \tag{0.2}$$

where the last approximation sign $\approx$ is due to the spatial dependence of the $\boldsymbol{c}$ on each other, and a pseudo probability is used. Then, from Bayes' formula, the following equation holds

$$\begin{aligned}&\log\left[\mathrm{P}(\theta \,|\, \mathbf{X}, \mathbf{Z})\right] \\ &= \log\left[\mathrm{P}(\mathbf{X}, \boldsymbol{f}, \boldsymbol{c} \,|\, \mathbf{L}, \mathbf{T}, \boldsymbol{\Lambda}, \boldsymbol{\gamma}, \boldsymbol{\mu}, \boldsymbol{\omega})\right] + \log\left[\mathrm{P}(\mathbf{L}, \boldsymbol{s} \,|\, \boldsymbol{p})\right] + \log\left[\pi(\mathbf{T})\pi(\boldsymbol{\Lambda})\pi(\boldsymbol{\gamma})\pi(\boldsymbol{\mu})\pi(\boldsymbol{\omega})\pi(\boldsymbol{p})\right] + C \\ &\approx \underbrace{\log \prod_{b,i} \mathrm{P}(\boldsymbol{x}_{bi}, \boldsymbol{f}_{bi}, c_{bi} \,|\, \hat{\boldsymbol{c}}_{\mathsf{N}_{bi}}, \mathbf{L}, \mathbf{T}_b, \boldsymbol{\Lambda}, \gamma_b, \boldsymbol{\mu}, \omega_{bi})}_{\widetilde{Q}_1} + \underbrace{\log\left[\mathrm{P}(\mathbf{L}, \boldsymbol{s} \,|\, \boldsymbol{p})\right]}_{Q_2} + \underbrace{\log\left[\pi(\mathbf{T})\pi(\boldsymbol{\Lambda})\pi(\boldsymbol{\gamma})\pi(\boldsymbol{\mu})\pi(\boldsymbol{\omega})\pi(\boldsymbol{p})\right]}_{Q_3} + C,\end{aligned} \tag{0.3}$$

where $\widetilde{Q}_1$ is a pseudo approximation of the complete-data log-likelihood corresponding to $(\mathbf{X}, \boldsymbol{f}, \boldsymbol{c})$ for the same reason as (0.2), $Q_2$ is the complete-data log-likelihood corresponding to the parameter $\mathbf{L}$ and $Q_3$ are joint priors. Finally, substituting (0.2) and (0.3) into $Q(\theta \,|\, \theta^{(t)})$, we get

$$\widetilde{Q}(\theta \,|\, \theta^{(t)}) = \mathbb{E}_{\widetilde{P}_1}[\widetilde{Q}_1] + \mathbb{E}_{P_2}[Q_2] + Q_3 + C, \tag{0.4}$$

which is a pseudo probability of $Q(\theta \,|\, \theta^{(t)})$ and where $\widetilde{P}_1 = P_{11}\widetilde{P}_{12}$.

In the maximization step, the block gradient descent algorithm is used to optimize $\widetilde{Q}(\theta \mid \theta^{(t)})$.

$$
\begin{aligned}
\mathbf{T}_b^{(t+1)} &= \arg\max_{T_b} \widetilde{Q}(T_b \mid \theta^{(t)}_{\setminus T_b}), && \text{for } b = 1, 2, \cdots, B, \\
\boldsymbol{\gamma}_b^{(t+1)} &= \arg\max_{\gamma_b} \widetilde{Q}(\boldsymbol{\gamma}_b \mid \theta^{(t)}_{\setminus \gamma_b}), && \text{for } b = 1, 2, \cdots, B, \\
\boldsymbol{\mu}_k^{(t+1)} &= \arg\max_{\mu_k} \widetilde{Q}(\boldsymbol{\mu}_k \mid \theta^{(t)}_{\setminus \boldsymbol{\mu}_k}), && \text{for } k = 1, 2, \cdots, q, \\
\omega_{bi}^{(t+1)} &= \arg\max_{\omega_{bi}} \widetilde{Q}(\omega_{bi} \mid \theta^{(t)}_{\setminus \omega_{bi}}), && \text{for } b = 1, 2, \cdots, B, \text{ and } i = 1, 2, \cdots, n_b, \\
\boldsymbol{\Lambda}^{(t+1)} &= \arg\max_{\Lambda} \widetilde{Q}(\boldsymbol{\Lambda} \mid \theta^{(t)}_{\setminus \boldsymbol{\Lambda}}), \\
p_j^{(t+1)} &= \arg\max_{p_j} \widetilde{Q}(p_j \mid \theta^{(t)}_{\setminus p_j}), && \text{for } j = 1, 2, \cdots, d, \\
L_{ij}^{(t+1)} &= \arg\max_{L_{ij}} \widetilde{Q}(L_{ij} \mid \theta^{(t)}_{\setminus L_{ij}}), && \text{for } i = 1, 2, \cdots, p, \text{ and } j = 1, 2, \cdots, d,
\end{aligned} \tag{0.5}
$$

where $\theta^{(t)}_{\setminus *}$ means all the parameters except for $*$.

#### Detailed calculations of Bayesian inference

**Calculation of Eq.** (0.1) First, based on Bayes' formula, we rewrite the log posterior $\log \mathrm{P}(\theta \mid \mathbf{X})$,

$$
\log [\mathrm{P}(\theta \mid \mathbf{X})] = \log [\mathrm{P}(\mathbf{X} \mid \theta)\pi(\theta)] - \log [\mathrm{P}(\mathbf{X})]\,. \tag{0.6}
$$

Then, due to the independence of $\mathrm{P}(\mathbf{X})$ with respect to $\theta$, the focus is mainly on the first term,

$$
\begin{aligned}
\log [\mathrm{P}(\mathbf{X} \mid \theta)\pi(\theta)] &= \log \left[ \int_z P(\mathbf{X}, \mathbf{Z} \mid \theta)\, \pi(\theta) \mathrm{d}z \right] \\
&= \log \left[ \int_z \mathrm{P}\Big(\mathbf{Z} \mid \mathbf{X}, \theta^{(t)}\Big) \frac{\mathrm{P}(\mathbf{X}, \mathbf{Z} \mid \theta)\, \pi(\theta)}{\mathrm{P}\big(\mathbf{Z} \mid \mathbf{X}, \theta^{(t)}\big)} \mathrm{d}z \right] \\
&\geqslant \int_z \mathrm{P}\Big(\mathbf{Z} \mid \mathbf{X}, \theta^{(t)}\Big) \log \left[ \frac{\mathrm{P}(\mathbf{X}, \mathbf{Z} \mid \theta)\, \pi(\theta)}{\mathrm{P}\big(\mathbf{Z} \mid \mathbf{X}, \theta^{(t)}\big)} \right] \mathrm{d}z \\
&= \mathbb{E}_{z \mid x, \theta^{(t)}} \log [\mathrm{P}(\theta \mid \mathbf{X}, \mathbf{Z})] + C(\theta^{(t)}),
\end{aligned} \tag{0.7}
$$

where the greater-than sign comes from Jensen's Inequality and $C(\theta^{(t)})$ is a constant with respect to $\theta$. Finally, by substituting (0.7) into (0.6) , we can obtain a lower bound on the log posterior $\log [\mathrm{P}(\theta \mid \mathbf{X})]$

$$
\log [\mathrm{P}(\theta \mid \mathbf{X})] \geqslant \mathbb{E}_{z \mid x, \theta^{(t)}} \log [\mathrm{P}(\theta \mid \mathbf{X}, \mathbf{Z})] + C,
$$

where $C = C(\theta^{(t)}) - \log [\mathrm{P}(\mathbf{X})]$ is also a constant.

**Calculation of Eq.** (0.2) The latent variable $\mathbf{Z}$ in our model consists of the low-dimensional batch-corrected factor $\boldsymbol{f}$, the cell type indicator $\boldsymbol{c}$, and the spike-and-slab distributional indicator $\boldsymbol{s}$. From the independence of $\boldsymbol{f}, \boldsymbol{c}$ and $\boldsymbol{s}$ under other random variables, the conditional distribution Eq. (0.2) could be computed separately, i.e., $P_{11}$, $\tilde{P}_{12}$ and $P_2$. Now, we compute $P_{11}$, $\tilde{P}_{12}$ and $P_2$ with $\theta^{(t)}$ shorted by $\theta$ for briefness,

$$
\begin{aligned}
P_{11} =& \prod_{b,i} \mathrm{P}(\boldsymbol{f}_{bi} \,|\, c_{bi}, \boldsymbol{x}_{bi}, \theta) = \prod_{b,i,k} \mathrm{P}(\boldsymbol{f}_{bi} \,|\, c_{bi} = k, \boldsymbol{x}_{bi}, \theta)^{\mathbb{I}(c_{bi}=k)} \\
=& \prod_{b,i,k} \exp\{\log\left[\mathrm{P}(\boldsymbol{x}_{bi} \,|\, \boldsymbol{f}_{bi}, \theta)\right] + \log\left[\mathrm{P}(\boldsymbol{f}_{bi} \,|\, c_{bi} = k, \theta)\right] - \log\left[\mathrm{P}(\boldsymbol{x}_{bi} \,|\, c_{bi} = k, \theta)\right]\}^{\mathbb{I}(c_{bi}=k)} \\
\propto& \prod_{b,i,k} \exp\left\{-\frac{1}{2}\Big[(\boldsymbol{x}_{bi} - \mathbf{L}\boldsymbol{f}_{bi} - \gamma_b)^\top \mathbf{T}_b (\boldsymbol{x}_{bi} - \mathbf{L}\boldsymbol{f}_{bi} - \gamma_b) + (\boldsymbol{f}_{bi} - \boldsymbol{\mu}_k)^\top \omega_{bi} \boldsymbol{\Lambda} (\boldsymbol{f}_{bi} - \boldsymbol{\mu}_k)\Big]\right\}^{\mathbb{I}(c_{bi}=k)} \\
\propto& \prod_{b,i,k} \exp\left\{-\frac{1}{2}\Big[\boldsymbol{f}_{bi}^\top (\mathbf{L}^\top \mathbf{T}_b \mathbf{L} + \omega_{bi}\boldsymbol{\Lambda})\boldsymbol{f}_{bi} - 2(\mathbf{L}^\top \mathbf{T}_b \widetilde{\boldsymbol{x}}_{bi} + \omega_{bi}\boldsymbol{\Lambda}\boldsymbol{\mu}_k)^\top \boldsymbol{f}_{bi}\Big]\right\}^{\mathbb{I}(c_{bi}=k)} \\
=& \prod_{b,i,k} \left[\mathcal{N}(\Phi_{bi}^{-1}\varphi_{bik}, \Phi_{bi}^{-1})\right]^{\mathbb{I}(c_{bi}=k)}, \\
\widetilde{P}_{12} =& \prod_{b,i} \mathrm{P}(c_{bi} \,|\, \hat{\boldsymbol{c}}_{\mathsf{N}_{bi}}, \boldsymbol{x}_{bi}, \theta) = \prod_{b,i,k} \left[\mathrm{P}(c_{bi} = k \,|\, \hat{\boldsymbol{c}}_{\mathsf{N}_{bi}}, \boldsymbol{x}_{bi}, \theta)\right]^{\mathbb{I}(c_{bi}=k)} \propto \prod_{b,i,k} \left[\mathrm{P}(\boldsymbol{x}_{bi}, c_{bi} = k \,|\, \hat{\boldsymbol{c}}_{\mathsf{N}_{bi}}, \theta)\right]^{\mathbb{I}(c_{bi}=k)} \\
=& \prod_{b,i,k} \left[\int_f \mathrm{P}(\boldsymbol{x}_{bi}, \boldsymbol{f}_{bi} \,|\, c_{bi} = k, \theta) \mathrm{d}f \times \mathrm{P}(c_{bi} = k \,|\, \hat{\boldsymbol{c}}_{\mathsf{N}_{bi}})\right]^{\mathbb{I}(c_{bi}=k)} \\
\propto& \prod_{b,i,k} \left\{\exp\Big[-\frac{1}{2}(\tilde{\boldsymbol{x}}_{bi} - \Psi_{bi}^{-1}\psi_{bik})^\top \Psi_{bi} (\tilde{\boldsymbol{x}}_{bi} - \Psi_{bi}^{-1}\psi_{bik}) + \eta_b \sum_{j \in \mathsf{N}_{bi}} \mathbb{I}(\hat{c}_j = k)\Big]\right\}^{\mathbb{I}(c_{bi}=k)} \\
=& \prod_{b,i,k} \left\{\exp\Big[-\frac{1}{2}(\tilde{\boldsymbol{x}}_{bi} - \mathbf{L}\boldsymbol{\mu}_k)^\top \Psi_{bi} (\tilde{\boldsymbol{x}}_{bi} - \mathbf{L}\boldsymbol{\mu}_k) + \eta_b \sum_{j \in \mathsf{N}_{bi}} \mathbb{I}(\hat{c}_j = k)\Big]\right\}^{\mathbb{I}(c_{bi}=k)} \\
\propto& \prod_{b,i,k} [S_{bik}]^{\mathbb{I}(c_{bi}=k)}, \\
P_2 =& \mathrm{P}(\boldsymbol{s} \,|\, \theta) = \prod_{i,j} \mathrm{P}(s_{ij} \,|\, L_{ij}, p_j) \propto \prod_{i,j} \mathrm{P}(L_{ij} \,|\, s_{ij}) \mathrm{P}(s_{ij} \,|\, p_j) \\
=& \Pi_{i,j} \left[\frac{p_j L_{ij}^2}{\lambda_1} |2\pi\lambda_1|^{-1/2} \exp\Big(-\frac{L_{ij}^2}{2\lambda_1}\Big)\right]^{s_{ij}} \left[(1 - p_j)|2\pi\lambda_0|^{-1/2} \exp\Big(-\frac{L_{ij}^2}{2\lambda_0}\Big)\right]^{1-s_{ij}} \\
=& \prod_{i,j} [h_{ij}]^{s_{ij}} [1 - h_{ij}]^{1-s_{ij}}.
\end{aligned}
$$

where $\propto$ means equal up to a constant, $\tilde{\boldsymbol{x}}_{bi} = \boldsymbol{x}_{bi} - \gamma_b$, $\Phi_{bi} = \mathbf{L}^\top \mathbf{T}_b \mathbf{L} + \omega_{bi}\boldsymbol{\Lambda}$, $\varphi_{bik} = \mathbf{L}^\top \mathbf{T}_b \tilde{\boldsymbol{x}}_{bi} + \omega_{bi}\boldsymbol{\Lambda}\boldsymbol{\mu}_k$, $\Psi_{bi} = \mathbf{T}_b - \mathbf{T}_b \mathbf{L} \Phi_{bi}^{-1} \mathbf{L}^\top \mathbf{T}_b$, $\psi_{bik} = \mathbf{T}_b \mathbf{L} \Phi_{bi}^{-1} \omega_{bi} \boldsymbol{\Lambda} \boldsymbol{\mu}_k$,

$$
h_{ij} = \frac{1}{1 + \frac{1-p_j}{p_j} \frac{\lambda_1}{L_{ij}^2} \sqrt{\frac{\lambda_1}{\lambda_0}} \exp\Big(-\frac{L_{ij}^2}{2}\Big(\frac{1}{\lambda_0} - \frac{1}{\lambda_1}\Big)\Big))},
$$

and

$$
S_{bik} \propto \exp\Big[-\frac{1}{2}(\tilde{\boldsymbol{x}}_{bi} - \mathbf{L}\boldsymbol{\mu}_k)^\top \Psi_{bi} (\tilde{\boldsymbol{x}}_{bi} - \mathbf{L}\boldsymbol{\mu}_k) + \eta_b \sum_{j \in \mathsf{N}_{bi}} \mathbb{I}(\hat{c}_j = k)\Big] \quad \text{such that} \quad \sum_k S_{bik} = 1.
$$

**Calculation of Eq. (0.3)** Based on Bayes' formula, the complete-data log posterior $\mathrm{P}(\theta \mid \mathbf{X}, \mathbf{Z})$ is split into three parts $\widetilde{Q}_1$, $Q_2$ and $Q_3$, which will be calculated in turn.

$$\begin{aligned}
\widetilde{Q}_1 &= \log\left[\prod_{b,i} \mathrm{P}(\boldsymbol{x}_{bi}, \boldsymbol{f}_{bi}, c_{bi} \mid \hat{\boldsymbol{c}}_{\mathsf{N}_{bi}}, \mathbf{L}, \mathbf{T}_b, \boldsymbol{\Lambda}, \boldsymbol{\mu}, \omega_{bi}, \boldsymbol{\gamma}_b)\right] \\
&= \log\left[\prod_{b,i,k} \Big\{\mathrm{P}(\boldsymbol{x}_{bi}, \boldsymbol{f}_{bi} \mid c_{bi} = k, \mathbf{L}, \mathbf{T}_b, \boldsymbol{\Lambda}, \boldsymbol{\mu}_{c_{bi}}, \omega_{bi}, \boldsymbol{\gamma}_{bi})\mathrm{P}(c_{bi} = k \mid \hat{\boldsymbol{c}}_{\mathsf{N}_{bi}})\Big\}^{\mathbb{I}(c_{bi}=k)}\right] \\
&= \sum_{b,i,k} \mathbb{I}(c_{bi} = k)\big\{ \log\left[\mathrm{P}(\boldsymbol{x}_{bi} \mid \boldsymbol{f}_{bi}, \mathbf{L}, \mathbf{T}_b, \boldsymbol{\gamma}_{bi})\right] + \log\left[\mathrm{P}(\boldsymbol{f}_{bi} \mid c_{bi} = k, \boldsymbol{\Lambda}, \boldsymbol{\mu}_k, \omega_{bi})\right] + \log\left[\mathrm{P}(c_{bi} = k \mid \hat{\boldsymbol{c}}_{\mathsf{N}_{bi}})\right]\big\} \\
&= \sum_{b,i}\sum_{k} \mathbb{I}(c_{bi} = k)\Big\{-\frac{1}{2}\operatorname{tr}\big[(\mathbf{L}^\top \mathbf{T}_b \mathbf{L} + \omega_{bi}\boldsymbol{\Lambda})\boldsymbol{f}_{bi}\boldsymbol{f}_{bi}^\top\big] + \big[(\boldsymbol{x}_{bi} - \boldsymbol{\gamma}_b)^\top \mathbf{T}_b \mathbf{L} + \omega_{bi}\boldsymbol{\mu}_k^\top \boldsymbol{\Lambda}\big]\boldsymbol{f}_{bi} \\
&\qquad + \frac{1}{2}\log|\mathbf{T}_b| + \frac{d}{2}\log|\omega_{bi}| + \frac{1}{2}\log|\boldsymbol{\Lambda}| - \frac{1}{2}(\boldsymbol{x}_{bi} - \boldsymbol{\gamma}_b)^\top \mathbf{T}_b(\boldsymbol{x}_{bi} - \boldsymbol{\gamma}_b) \\
&\qquad - \frac{1}{2}\omega_{bi}\boldsymbol{\mu}_k^\top \boldsymbol{\Lambda}\boldsymbol{\mu}_k + \eta_b \sum_{j \in \mathsf{N}_{bi}} \mathbb{I}(\hat{c}_j = k)\Big\} + C,
\end{aligned}$$

$$\begin{aligned}
Q_2 &= \log\left[\mathrm{P}\big(\mathbf{L}, \boldsymbol{s} \mid \boldsymbol{p}\big)\right] = \log\left[\prod_{i,j} \mathrm{P}\big(L_{ij}, s_{ij} \mid p_j\big)\right] \\
&= \sum_{i,j} \log\left[\prod_{s \in \{0,1\}} \Big[\mathrm{P}\big(s_{ij} = s \mid p_j\big)\mathrm{P}\big(L_{ij} \mid s_{ij} = s\big)\Big]^{\mathbb{I}(s_{ij}=s)}\right] \\
&= \sum_{i,j} \Big\{\mathbb{I}(s_{ij} = 1)\Big[\log p_j + \log \mathrm{P}\big(L_{ij} \mid s_{ij} = 1\big)\Big] + \mathbb{I}(s_{ij} = 0)\Big[\log(1 - p_j) + \log \mathrm{P}\big(L_{ij} \mid s_{ij} = 0\big)\Big]\Big\} \\
&= \sum_{i,j} \Big\{\mathbb{I}(s_{ij} = 1)\Big[\log p_j + \log \frac{L_{ij}^2}{\lambda_1} - \frac{L_{ij}^2}{2\lambda_1}\Big] + \mathbb{I}(s_{ij} = 0)\Big[\log(1 - p_j) - \frac{L_{ij}^2}{2\lambda_0}\Big]\Big\} + C,
\end{aligned}$$

$$\begin{aligned}
Q_3 &= \log\left[\pi(\boldsymbol{\gamma})\pi(\boldsymbol{\mu})\pi(\boldsymbol{\omega})\pi(\boldsymbol{p})\pi(\mathbf{T})\pi(\boldsymbol{\Lambda})\right] \\
&= \sum_{b}\Big(-\frac{1}{2}\|\boldsymbol{\gamma}_b\|_2^2\Big) + \sum_{k}\Big[-\frac{1}{2}(\boldsymbol{\mu}_k - \boldsymbol{\mu}_\mu)^\top \boldsymbol{\Sigma}_\mu^{-1}(\boldsymbol{\mu}_k - \boldsymbol{\mu}_\mu)\Big] + \sum_{b,i}\Big[\Big(\frac{\nu_\omega}{2} - 1\Big)\log \omega_{bi} - \frac{\nu_\omega}{2}\omega_{bi}\Big] \\
&\quad + \sum_{b}\Big[\Big(\frac{\nu_\tau}{2} - 1\Big)\log|\mathbf{T}_b| - \frac{\nu_\tau}{2}\operatorname{tr}(\mathbf{T}_b)\Big] + \Big[\frac{n_\Lambda - d - 1}{2}\log|\boldsymbol{\Lambda}| - \frac{1}{2}\operatorname{tr}\big(\boldsymbol{\Sigma}_\Lambda^{-1}\boldsymbol{\Lambda}\big)\Big] \\
&\quad + \sum_{j}\Big[\Big(\frac{\alpha_p}{j} - 1\Big)\log p_j + (\beta_p - 1)\log(1 - p_j)\Big].
\end{aligned}$$

**Calculation of Eq. (0.4)** Substituting Eq. (0.2) into Eq. (0.3) gives Eq. (0.4).

$$\begin{aligned}
\mathbb{E}_{\tilde{P}_1}[\tilde{Q}_1] = \sum_{b,i}\sum_{k} S_{bik}\Big\{&\frac{1}{2}\log|\mathbf{T}_b| + \frac{d}{2}\log|\omega_{bi}| + \frac{1}{2}\log|\boldsymbol{\Lambda}| - \frac{1}{2}(\boldsymbol{x}_{bi} - \boldsymbol{\gamma}_b)^\top \mathbf{T}_b(\boldsymbol{x}_{bi} - \boldsymbol{\gamma}_b) \\
&- \frac{1}{2}\boldsymbol{\mu}_k^\top \omega_{bi}\boldsymbol{\Lambda}\boldsymbol{\mu}_k + \eta_b \sum_{j \in \mathsf{N}_{bi}} \mathbb{I}(\hat{c}_j = k) - \frac{1}{2}\operatorname{tr}\Big[(\mathbf{L}^\top \mathbf{T}_b \mathbf{L} + \omega_{bi}\boldsymbol{\Lambda})\mathbb{E}_{P_{11} \mid c_{bi}=k}\Big(\boldsymbol{f}_{bi}\boldsymbol{f}_{bi}^\top\Big)\Big] \\
&+ \Big[(\boldsymbol{x}_{bi} - \boldsymbol{\gamma}_b)^\top \mathbf{T}_b \mathbf{L} + \boldsymbol{\mu}_k^\top \omega_{bi}\boldsymbol{\Lambda}\Big]\mathbb{E}_{P_{11} \mid c_{bi}=k}\Big(\boldsymbol{f}_{bi}\Big)\Big\} + C,
\end{aligned}$$

$$\mathbb{E}_{P_2}[Q_2] = \sum_{i,j}\Big\{h_{ij}\Big[\log p_j + \log\frac{L_{ij}^2}{\lambda_1} - \frac{L_{ij}^2}{2\lambda_1}\Big] + (1-h_{ij})\Big[\log(1-p_j) - \frac{L_{ij}^2}{2\lambda_0}\Big]\Big\} + C.$$

All expectations used in calculation of Eq. (0.4) are

$$\begin{aligned}
\mathbb{E}_{P_2}[\mathbb{I}(s_{ij}=1)] &= h_{ij}, \\
\mathbb{E}_{P_{11}\,|\,c_{bi}=k}[\boldsymbol{f}_{bi}] &\overset{\text{def}}{=} \mathbb{E}_{bik}[\boldsymbol{f}_{bi}] = \Phi_b^{-1}\varphi_{bik}, \\
\mathbb{E}_{P_{11}\,|\,c_{bi}=k}\Big[\boldsymbol{f}_{bi}\boldsymbol{f}_{bi}^\top\Big] &\overset{\text{def}}{=} \mathbb{E}_{bik}\Big[\boldsymbol{f}_{bi}\boldsymbol{f}_{bi}^\top\Big] = \Phi_{bi}^{-1} + \Phi_{bi}^{-1}\varphi_{bik}\varphi_{bik}^\top\Phi_{bi}^{-1}.
\end{aligned}$$

And updating subtype indicators $\boldsymbol{c}$ uses

$$\hat{c}_{bi}^{(t+1)} = \arg\max_k S_{bik} = \arg\max_k \exp\Big[-\frac{1}{2}(\tilde{\boldsymbol{x}}_{bi} - \mathbf{L}\boldsymbol{\mu}_k)^\top\Psi_{bi}(\tilde{\boldsymbol{x}}_{bi} - \mathbf{L}\boldsymbol{\mu}_k) + \eta_b \sum_{j\in \mathsf{N}_{bi}} \mathbb{I}(\hat{c}_j = k)\Big].$$

**Calculation of Eq. (0.5)** To obtain Eq. (0.5), we will derive the partial derivative of each variable under all the others fixed, then obtain the following

$$\begin{aligned}
\{\mathbf{T}_b^{-1}\}^{(t+1)} =& \frac{1}{n_b + \nu_\tau - 2}\text{diag}\Big\{\sum_{i\in \mathsf{B}_b}\Big[\tilde{\boldsymbol{x}}_{bi}\tilde{\boldsymbol{x}}_{bi}^\top + \mathbf{L}\Big(\sum_k S_{bik}\mathbb{E}_{bik}\Big[\boldsymbol{f}_{bi}\boldsymbol{f}_{bi}{}^\top\Big]\Big)\mathbf{L}^\top \\
& -\tilde{\boldsymbol{x}}_{bi}\Big(\sum_k S_{bik}\mathbb{E}_{bik}[\boldsymbol{f}_{bi}]\Big)^\top\mathbf{L}^\top - \mathbf{L}\Big(\sum_k S_{bik}\mathbb{E}_{bik}[\boldsymbol{f}_{bi}]\Big)\tilde{\boldsymbol{x}}_{bi}^\top\Big] + \nu_\tau\mathbf{I}_p\Big\}, \\
\gamma_b^{(t+1)} =& (n_b\mathbf{T}_b + \mathbf{I}_p)^{-1}\sum_{i\in \mathsf{B}_b}\mathbf{T}_b\Big(\boldsymbol{x}_{bi} - \mathbf{L}\sum_k S_{bik}\mathbb{E}_{bik}[\boldsymbol{f}_{bi}]\Big), \\
\boldsymbol{\mu}_k^{(t+1)} =& \Big(\sum_{b,i} S_{bik}\omega_{bi}\boldsymbol{\Lambda} + \boldsymbol{\Sigma}_\mu^{-1}\Big)^{-1}\Big(\sum_{b,i} S_{bik}\omega_{bi}\boldsymbol{\Lambda}\mathbb{E}_{bik}[\boldsymbol{f}_{bi}] + \boldsymbol{\Sigma}_\mu^{-1}\boldsymbol{\mu}_\mu\Big), \\
\omega_{bi}^{(t+1)} =& \frac{d + \nu_\omega - 2}{\sum_k S_{bik}\Big(\boldsymbol{\mu}_k^\top\boldsymbol{\Lambda}\boldsymbol{\mu}_k + \text{tr}\big(\boldsymbol{\Lambda}\mathbb{E}_{bik}\big[\boldsymbol{f}_{bi}\boldsymbol{f}_{bi}{}^\top\big]\big) - 2\boldsymbol{\mu}_k^\top\boldsymbol{\Lambda}\mathbb{E}_{bik}[\boldsymbol{f}_{bi}]\Big) + \nu_\omega}, \\
\{\boldsymbol{\Lambda}^{-1}\}^{(t+1)} =& \frac{1}{n + n_\Lambda - d - 1}\Big\{\sum_{b,i,k}\omega_{bi}S_{bik}\Big(\mathbb{E}_{bik}\Big[\boldsymbol{f}_{bi}\boldsymbol{f}_{bi}{}^\top\Big] + \boldsymbol{\mu}_k\boldsymbol{\mu}_k^\top - \boldsymbol{\mu}_k\mathbb{E}_{bik}[\boldsymbol{f}_{bi}]^\top - \mathbb{E}_{bik}[\boldsymbol{f}_{bi}]\,\boldsymbol{\mu}^\top\Big) + \boldsymbol{\Sigma}_\Lambda^{-1}\Big\}, \\
p_j^{(t+1)} =& \frac{\sum_i h_{ij} + \alpha_{p_j}/j - 1}{p + \alpha_{p_j} + \beta_p - 2}, \\
\frac{\partial\widetilde{Q}(\theta\mid\theta^{(t)})}{\partial L_{ij}} =& \frac{1}{L_{ij}}\Big\{\underbrace{-\Big[\frac{h_{ij}}{\lambda_1} + \frac{1-h_{ij}}{\lambda_0} + \sum_{b,s}\tau_{bi}\Big(\sum_k S_{bsk}\mathbb{E}_{bik}\big[f_{bsj}{}^2\big]\Big)\Big]}_{a_{ij}} L_{ij}^2 + \underbrace{2h_{ij}}_{c_{ij}} \\
& + \underbrace{\sum_{b,s}\Big[\tau_{bi}\tilde{\boldsymbol{x}}_{bsi}\Big(\sum_k S_{bsk}\mathbb{E}_{bik}[f_{bsj}]\Big) - \sum_{u\neq j}\tau_{bi}L_{iu}\sum_k S_{bsk}\Big(\mathbb{E}_{bik}\big(f_{bsu}f_{bsj}\big)\Big)\Big]}_{b_{ij}} L_{ij}\Big\}.
\end{aligned}$$

Define

$$\overline{L}_{ij} = \frac{-b_{ij} + \sqrt{b_{ij}^2 - 4a_{ij}c_{ij}}}{2a_{ij}}, \qquad \underline{L}_{ij} = \frac{-b_{ij} - \sqrt{b_{ij}^2 - 4a_{ij}c_{ij}}}{2a_{ij}},$$

then from Lemma 1 in Avalos-Pacheco *et al.* (2022)[2], we have

$$\widehat{L}_{ij} = \mathbb{I}(b_{ij} < 0)\, \overline{L}_{ij} + \mathbb{I}(b_{ij} > 0)\, \underline{L}_{ij}.$$

To make our paper self-contained, the lemma is given below.

**Lemma 0.2** (Avalos-Pacheco *et al.* (2022)[2]). *Let $f(x) = ax^2 + bx + c\log\big(x^2\big)$, where $a < 0$ and $c > 0$, with notation $\Delta = b^2 - 16ac$, we have*

$$\widehat{x} = \arg\max_x f(x) = \begin{cases} 0 & \textit{if } \Delta < 0, \\ \frac{-b+\sqrt{\Delta}}{4a} & \textit{if } \Delta \geqslant 0 \textit{ and } b < 0, \\ \frac{-b-\sqrt{\Delta}}{4a} & \textit{if } \Delta \geqslant 0 \textit{ and } b > 0, \\ \pm\sqrt{\frac{-c}{a}} & \textit{if } \Delta \geqslant 0 \textit{ and } b > 0. \end{cases}$$

*Proof.* Taking derivative of $f(x)$ with respect to $x$ is

$$\frac{\mathrm{d}f(x)}{\mathrm{x}} = 2ax + b + \frac{2c}{x} = \frac{1}{x}\big(2ax^2 + bx + 2c\big)\,,$$

with corresponding discriminant denoted by

$$\Delta = b^2 - 16ac \geqslant 0,$$

wherein $\Delta$ being always positive is from $a < 0$ and $c > 0$. Then roots formula of quadratic equation gives

$$\bar{x} = \frac{-b + \sqrt{\Delta}}{4a} \leqslant 0, \qquad \underline{x} = \frac{-b - \sqrt{\Delta}}{4a} \geqslant 0.$$

To check which of $\bar{x}$ and $\underline{x}$ is maximizer, their functional values are compared

$$f(\bar{x}) - f(\underline{x}) = \frac{b\sqrt{\Delta}}{4a} + c\log\left[\left(\frac{-b+\sqrt{\Delta}}{-b-\sqrt{\Delta}}\right)^2\right].$$

- If $b > 0$, with $a < 0$, we have $b\sqrt{\Delta}/a < 0$ and $\mid -b + \sqrt{\Delta} \mid < \mid -b - \sqrt{\Delta} \mid$, then $f(\bar{x}) < f(\underline{x})$.
- If $b < 0$, with $a < 0$, we have $b\sqrt{\Delta}/a > 0$ and $\mid -b + \sqrt{\Delta} \mid > \mid -b - \sqrt{\Delta} \mid$, then $f(\bar{x}) > f(\underline{x})$.
- If $b = 0$, we have $b\sqrt{\Delta}/a = 0$ and $\mid -b + \sqrt{\Delta} \mid = \mid -b - \sqrt{\Delta} \mid$, then $f(\bar{x}) = f(\underline{x})$.

□

#### Compared methods

We compared STADIA with three competing data integration methods, including PRECAST, fastMNN, and Harmony. For fairness of comparison, all methods adopt the same data pre-processing procedure, described in the main text. And the implementation of all compared methods were followed the authors' instructions.

- **PRECAST[3].** First `GetAssayData` and *$* functions from the R package Seurat are used to extract gene expression profiles and coordinates for all spots. Then `ICM.EM` function from the R package PRECAST (v1.6) with low dimensionality parameter $q$ equal to 35. Finally select the final result using the function `selectModel` in the R package PRECAST.

- **fastMNN[4].** FastMNN is a fast version of MNN following principal component analysis (PCA). Based on the selected HVGs, we used the `RunFastMNN` function in the R package SeuratWrappers (v0.3.1), adapting `fastMNN` in the R package batchelor, with the top 35 PCs and all other parameters by default.

- **Harmony[5].** First get the top 35 PCs with the function `RunPCA` in the R package Seurat (v4.3.0) and then do Harmony with the function `RunHarmony` in the R package harmony (v0.1.1) with all default parameters.

#### Evaluation metric

- **Adjusted Rand Index (ARI)[6]**, adapting the Rand Index (RI), measures the similarity between two partitions, defined as

$$\text{ARI} = \frac{\sum_{ij}\binom{n_{ij}}{2} - \left[\sum_i \binom{a_i}{2}\sum_j \binom{b_j}{2}\right] / \binom{n}{2}}{\frac{1}{2}\left[\sum_i \binom{a_i}{2} + \sum_j \binom{b_j}{2}\right] - \left[\sum_i \binom{a_i}{2} + \sum_j \binom{b_j}{2}\right]\binom{n}{2}},$$

  where $a$ and $b$ are two partitions, $n$ is the total number of nodes, $a_i$ and $b_j$ are the number of nodes in particular partitions, and $n_{ij}$ is the number of cells that appear simultaneously in the $i$th cluster of partition $a$ and the $j$th cluster of partition $b$. ARI is a symmetric measure that ranges from $0$ to $1$, and higher value indicates higher similarity. We compute ARI to compare the clustering result of the integrated data with the predefined cell types.

- **Normalized Mutual Information (NMI)[7]**, is the normalization of the Mutual Information (MI) used to measure clustering accuracy. NMI is defined as

$$\text{NMI} = 2 \times \frac{\sum_{ij}\frac{n_{ij}}{n}\log\left(\frac{n \times n_{ij}}{a_i \times b_j}\right)}{\sum_i \frac{a_i}{n}\log\left(\frac{n}{a_i}\right) + \sum_j \frac{b_j}{n}\log\left(\frac{n}{b_j}\right)},$$

  where the notation is the same as that in ARI. NMI ranges in $[0, 1]$ and higher value also indicates higher similarity between the clustering result and the true cell types.

- **Local Inverse Simpson's Index of Integration (LISI)[5]** is used to evaluate the quality of mixing after integration. Based on local neighbors chosen on a preselected perplexity, LISI calculates the effective number of different batches in the local neighborhood of each cell for *iLISI* score, and of different cell types for *cLISI* score. From a perspective of application, the higher value of iLISI and lower value of cLISI mean better mixing.

### Supplementary Figures

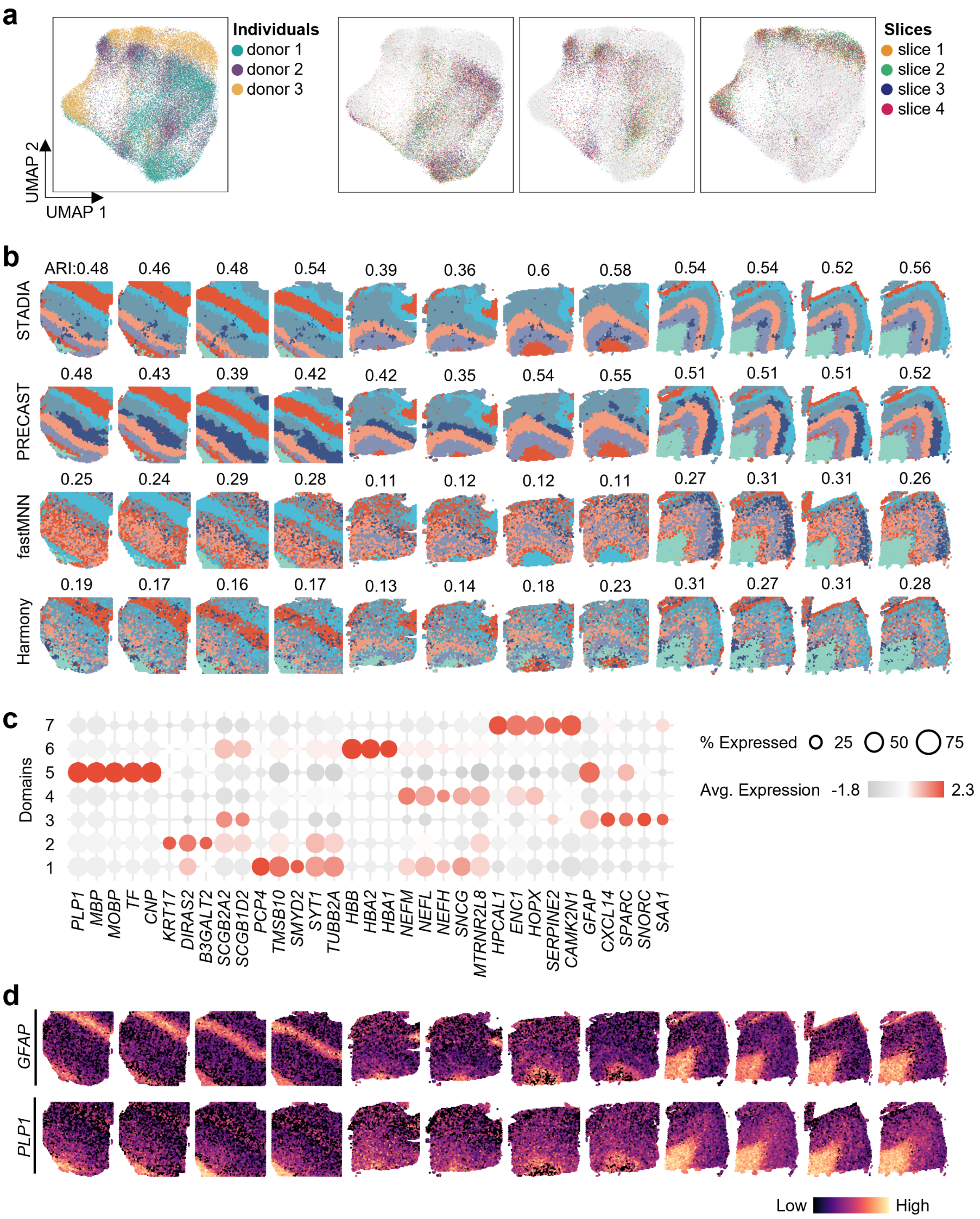


**Fig. S1.** Integration of 12 slices of DLPFC dataset. **a.** Embedded UMAP plots for original raw data colored by different donors (left) and slices per donor (left). **b.** Visualization of the spatial domains for the four methods with the corresponding ARI marked on each slice. **c.** Pearson correlations between the domains identified by STADIA. **d.** Dot plot of the top 5 marker genes for each domain identified by STADIA. **e.** Visualization of the expression of markers for layer 1 and WM in spatial context.

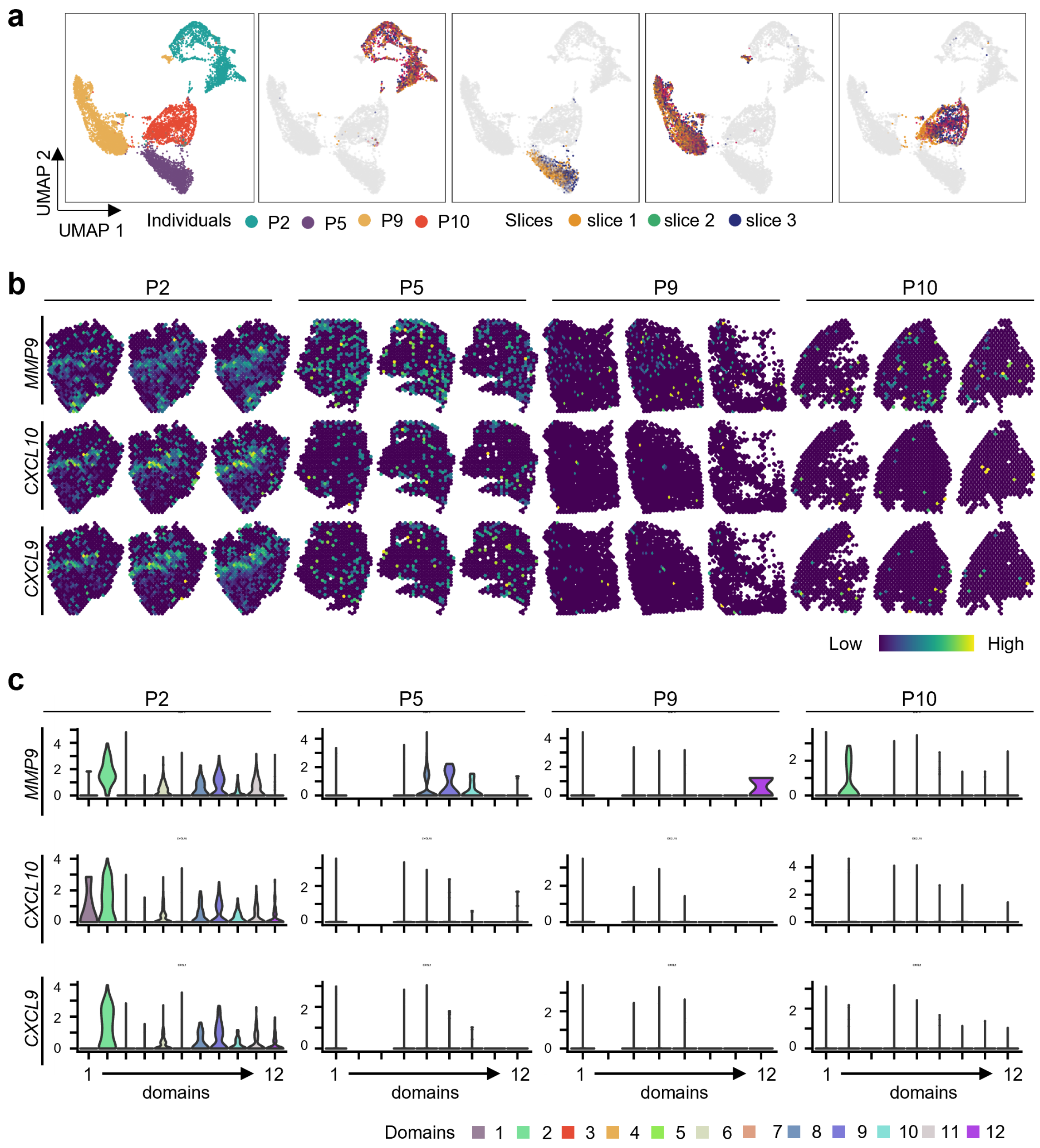


**Fig. S2.** Integration of 12 slices of cSCC dataset. **a.** Embedded UMAP plots for original raw data colored by different patients (left) and slices per patient (left). **b.** Visualization of the expression of markers *MMP9*, *CXCL10*, *CXCL9* for domain 2 in spatial context. **c.** Violin plot of the expression of genes *MMP9*, *CXCL10*, *CXCL9*.

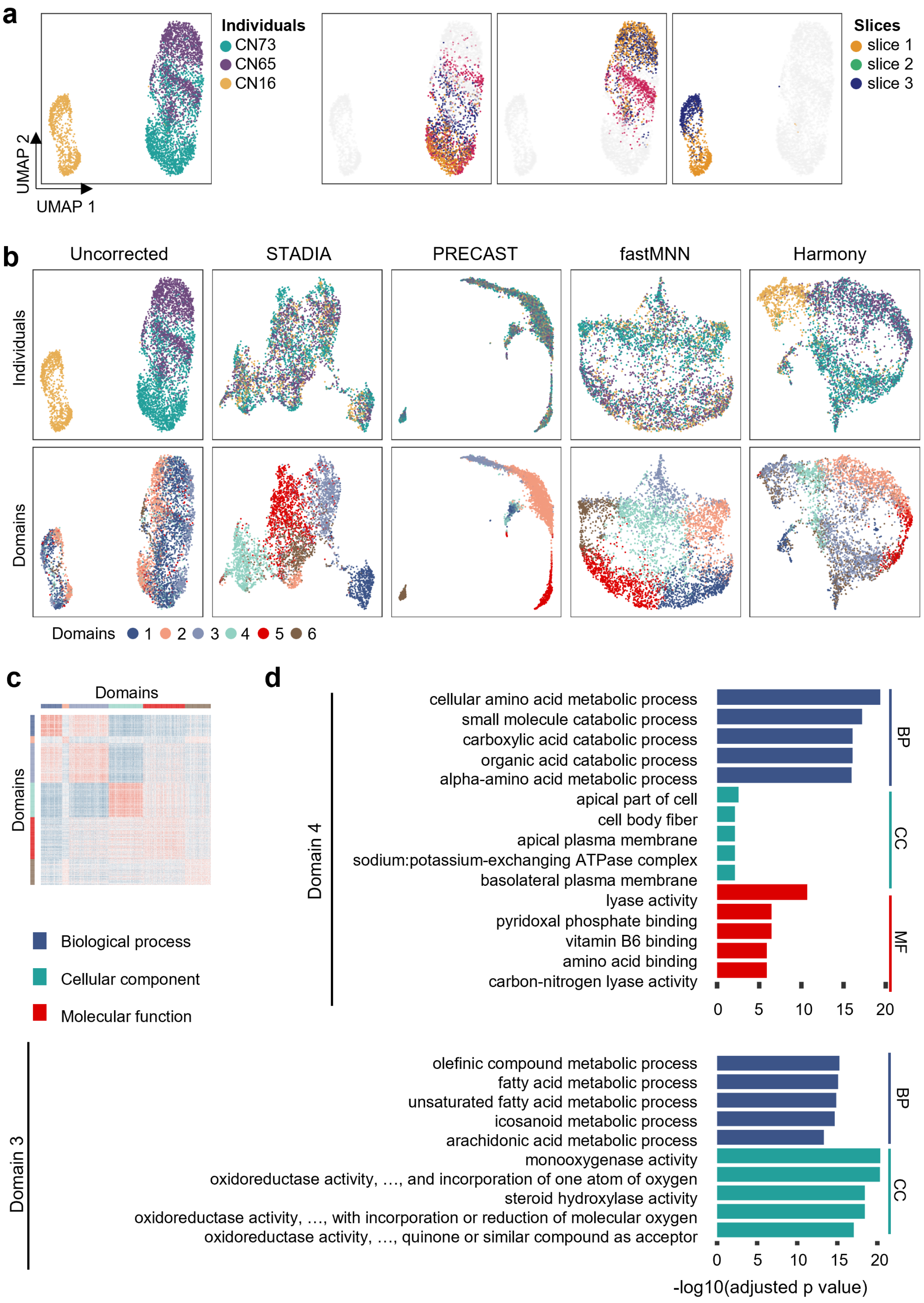
a
Individuals
CN73
CN65
CN16
UMAP 2
UMAP 1
Slices
slice 1
slice 2
slice 3
b
Uncorrected
STADIA
PRECAST
fastMNN
Harmony
Individuals
Domains
Domains 1 2 3 4 5 6
c
Domains
Domains
Biological process
Cellular component
Molecular function
d
Domain 4
cellular amino acid metabolic process
small molecule catabolic process
carboxylic acid catabolic process
organic acid catabolic process
alpha-amino acid metabolic process
apical part of cell
cell body fiber
apical plasma membrane
sodium:potassium-exchanging ATPase complex
basolateral plasma membrane
lyase activity
pyridoxal phosphate binding
vitamin B6 binding
amino acid binding
carbon-nitrogen lyase activity
BP
CC
MF
0 5 10 15 20
Domain 3
olefinic compound metabolic process
fatty acid metabolic process
unsaturated fatty acid metabolic process
icosanoid metabolic process
arachidonic acid metabolic process
monooxygenase activity
oxidoreductase activity, …, and incorporation of one atom of oxygen
steroid hydroxylase activity
oxidoreductase activity, …, with incorporation or reduction of molecular oxygen
oxidoreductase activity, …, quinone or similar compound as acceptor
BP
CC
0 5 10 15 20
-log10(adjusted p value)

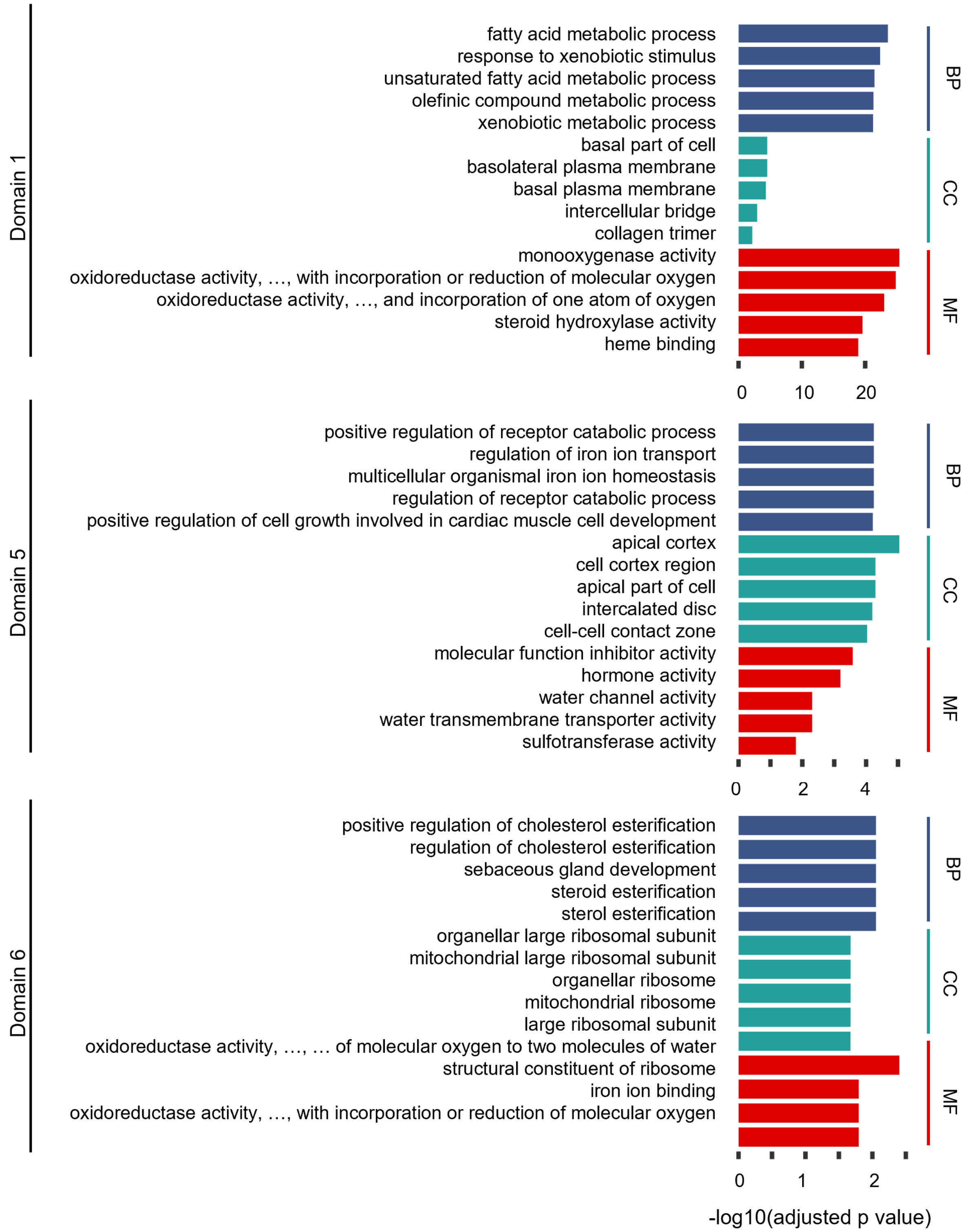


**Fig. S3.** Integration of 12 slices of cSCC dataset. **a.** Embedded UMAP plots for original raw data colored by different mice (left) and slices per mouse (left). **b.** Embedded UMAP plots for the four methods colored by mice (top) and spatial domains (bottom). **c.** Pearson correlations between the domains identified by STADIA. **d.** The top 5 most significant GO terms for all domains except for doamin 2, which is also in **Fig. 4**.

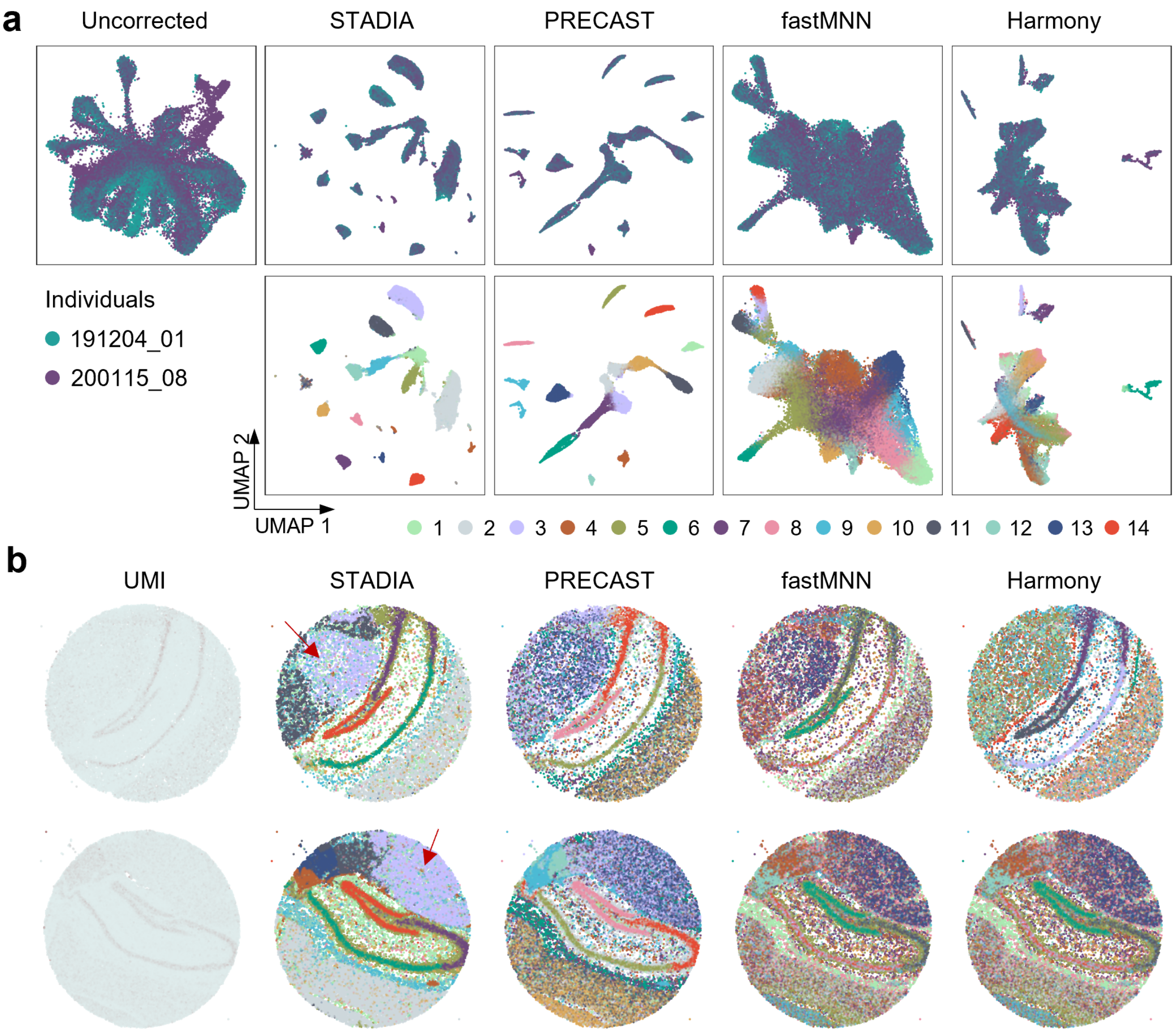


**Fig. S4.** Integration of two slices of the hippocampal dataset profiled by slide-seqV2. **a.** Embedded UMAP plots for the four methods colored by slices (top) and spatial domains (bottom). **b.** Visualization of spatial domains identified by the four methods in spatial context.

#### Supplementary Table

**Table S1.** Summary of all ST data used in this study.

| Platform | Species | Tissue | Slices | Spots | Related Figures | Reference |
|---|---|---|---|---|---|---|
| 10x Visium | Human | Dorsolateral prefrontal cortex (DLPFC) | 151507 | 4226 | Fig. 2 Fig. S1 | [8] |
| | | | 151508 | 4384 | | |
| | | | 151509 | 4789 | | |
| | | | 151510 | 4634 | | |
| | | | 151669 | 3661 | | |
| | | | 151670 | 3498 | | |
| | | | 151671 | 4110 | | |
| | | | 151672 | 4015 | | |
| | | | 151673 | 3639 | | |
| | | | 151674 | 3673 | | |
| | | | 151675 | 3592 | | |
| | | | 151676 | 3460 | | |
| | Mouse | Brain | Anterior | 2696 | Fig. 3 | 10x Visium demo |
| | | | Posterior | 3353 | | |
| ST | Human | Cutaneous squamous cell carcinoma (cSCC) | P2_rep1 | 666 | Fig. 5 Fig. S3 | [9] |
| | | | P2_rep2 | 645 | | |
| | | | P2_rep3 | 638 | | |
| | | | P5_rep1 | 584 | | |
| | | | P5_rep2 | 517 | | |
| | | | P5_rep3 | 517 | | |
| | | | P9_rep1 | 1125 | | |
| | | | P9_rep2 | 1035 | | |
| | | | P9_rep3 | 828 | | |
| | | | P10_rep1 | 545 | | |
| | | | P10_rep2 | 619 | | |
| | | | P10_rep3 | 460 | | |
| | Mouse | Liver | CN73_C1 | 673 | Fig. 4 Fig. S2 | [10] |
| | | | CN73_D1 | 684 | | |
| | | | CN73_E2 | 650 | | |
| | | | CN65_D1 | 663 | | |
| | | | CN65_D2 | 629 | | |
| | | | CN65_E1 | 590 | | |
| | | | CN16_D2 | 487 | | |
| | | | CN16_E2 | 487 | | |
| Slide-seqV2 | Mouse | Hippocampus | Puck_191204_01 | 34199 | Fig. 6 | [11] |
| | | | Puck_200115_08 | 53208 | | |

**Table S2.** The Allen Reference Atlas of Mouse Brain.

| Related figures | ABA ID | ABA URL |
|---|---|---|
| Fig. 3a | 100883818 | http://atlas.brain-map.org/atlas?atlas=2&plate=100883818 |
| Fig. 6a | 100960084 | http://atlas.brain-map.org/atlas?atlas=1&plate=100960084 |

**Table S3.** Parameters Used in Experiments.

| Dataset | d | K | eta |
|---|---|---|---|
| Human dorsolateral prefrontal cortex (DLPFC) | 35 | 7 | 0.23 |
| Mouse brain | 35 | 35 | 0.15 |
| Human cutaneous squamous cell carcinoma (cSCC) | 35 | 12 | 0.15 |
| Mouse liver | 35 | 6 | 0.15 |
| Mouse Hippocampus | 35 | 14 | 0.23 |

#### References


1. Liu, C. & Rubin, D. B. ML estimation of the t distribution using EM and its extensions, ECM and ECME. *Statistica Sinica* **5,** 19–39 (1995).
2. Avalos-Pacheco, A., Rossell, D. & Savage, R. S. Heterogeneous large datasets integration using Bayesian factor regression. *Bayesian Analysis* **17,** 33–66 (2022).
3. Liu, W. *et al.* Probabilistic embedding, clustering, and alignment for integrating spatial transcriptomics data with PRECAST. *Nature Communications* **14,** 296 (2023).
4. Lun, A. Further MNN algorithm development. https://MarioniLab.github.io/FurtherMNN2018/theory/description.html. (2019).
5. Korsunsky, I. *et al.* Fast, sensitive and accurate integration of single-cell data with Harmony. *Nature Methods* **16,** 1289–1296 (2019).
6. Hubert, L. & Arabie, P. Comparing partitions. *Journal of Classification* **2,** 193–218 (1985).
7. Shannon, C. E. A mathematical theory of communication. *The Bell System Technical Journal* **27,** 379–423 (1948).
8. Maynard, K. R. *et al.* Transcriptome-scale spatial gene expression in the human dorsolateral prefrontal cortex. *Nature Neuroscience* **24,** 425–436 (2021).
9. Ji, A. L. *et al.* Multimodal analysis of composition and spatial architecture in human squamous cell carcinoma. *Cell* **182,** 497–514 (2020).
10. Hildebrandt, F. *et al.* Spatial Transcriptomics to define transcriptional patterns of zonation and structural components in the mouse liver. *Nature Communications* **12,** 7046 (2021).
11. Stickels, R. R. *et al.* Highly sensitive spatial transcriptomics at near-cellular resolution with Slide-seqV2. *Nature Biotechnology* **39,** 313–319 (2021).
